# NDR1/2 kinases regulate membrane trafficking, enable efficient autophagy and prevent neurodegeneration

**DOI:** 10.1101/2022.03.28.486054

**Authors:** Flavia Roșianu, Simeon R Mihaylov, Noreen Eder, Antonie Martiniuc, Suzanne Claxton, Helen R Flynn, Shamsinar Jalal, Marie-Charlotte Domart, Lucy Collinson, Mark Skehel, Ambrosius P Snijders, Matthias Krause, Sharon A Tooze, Sila K Ultanir

## Abstract

Autophagy is essential for neuronal development and its deregulation contributes to neurodegenerative diseases. NDR1 and NDR2 are highly conserved kinases implicated in neuronal development, mitochondrial health and autophagy, but how they affect mammalian brain development *in vivo* is not known. Using single and double *Ndr1/2* knockout mouse models we show that, dual, but not individual loss of *Ndr1/2* in neurons causes neurodegeneration during brain development, but also in adult mice. Proteomic and phosphoproteomic comparisons between *Ndr1/2* knockout and control brains revealed novel kinase substrates and indicated that endocytosis is significantly affected in the absence of NDR1/2. We validated the endocytic protein, Raph1/Lpd1 as a novel NDR1/2 substrate and showed that both NDR1/2 and Raph1 are critical for endocytosis and membrane recycling. In NDR1/2 knockout brains, we observed prominent accumulation of transferrin receptor, p62 and ubiquitinated proteins, indicative of a major impairment of protein homeostasis. Furthermore, the levels of LC3-positive autophagosomes were reduced in knockout neurons, implying that reduced autophagy efficiency mediates p62 accumulation and neurotoxicity. Mechanistically, pronounced mislocalisation of the transmembrane autophagy protein ATG9A at the neuronal periphery, impaired axonal ATG9A trafficking and increased ATG9A surface levels further confirm defects in membrane trafficking and could underlie the impairment in autophagy. We provide novel insight into the roles of NDR1/2 kinases in maintaining neuronal health.

**Highlights:**

- Dual neuronal *Ndr1* and *Ndr2* knockout during development or in adult mice causes neurodegeneration.
- Phosphoproteomics comparison of *Ndr1/2* knockouts with control littermates shows endocytosis and membrane trafficking to be affected and reveals novel substrates.
- Raph1/Lamellipodin is a novel NDR1/2 substrate that is required for TfR endocytosis.
- *Ndr1/2* knockout brains exhibit a severe defect in ubiquitinated protein clearance and reduced autophagy.
- NDR1/2 and Raph1 are required for the trafficking of the only transmembrane autophagy protein, ATG9A.

## Introduction

Macroautophagy (hereafter referred to as autophagy) is a degradation process for cytoplasmic organelles and proteins (Klionsky *et al*, 2010; Mizushima & Komatsu, 2011; Ohsumi, 2001; Yamamoto & Yue, 2014; Yorimitsu & Klionsky, 2005; Yu *et al*, 2018). Due to its essential role in maintaining neuronal protein and organelle homeostasis, via a constant turnover of these components, constitutive autophagy is indispensable for neuronal survival (Azarnia Tehran *et al*, 2018; Kulkarni *et al*, 2018; Menzies *et al*, 2017; Stavoe & Holzbaur, 2019). Neuronal autophagosomes form predominantly in the axons and presynaptic compartments and are trafficked to the soma, maturing along the way and eventually fusing with fully competent lysosomes in the cell body (Maday *et al*, 2014; Murdoch *et al*, 2016; Soukup *et al*, 2016; Stavoe & Holzbaur, 2019). Impairment of autophagy in brain-specific *Atg7* or *Atg5* knockout mice leads to neurodegeneration (Hara *et al*, 2006; Komatsu *et al*, 2006) and impairment of autophagy may also underlie human neurodegenerative diseases (Harris & Rubinsztein, 2011; Menzies *et al*., 2017; Wong & Holzbaur, 2015). Autophagy has long been considered an essential mechanism for neuronal homeostasis that is needed to prevent neurodegeneration and enhancing autophagy is a putative therapeutic avenue for neurodegenerative disorders with misfolded protein accumulations (Djajadikerta *et al*, 2020; Karabiyik *et al*, 2017; Menzies *et al*., 2017). ATG9A is the only transmembrane autophagy component and functions at the early steps of autophagosome formation (Webber & Tooze, 2010; Young *et al*, 2006). In mammalian cells, ATG9A transiently interacts with and contributes to phagophore formation (Judith *et al*, 2019; Orsi *et al*, 2012). ATG9A cycles between Golgi, recycling endosomes and plasma membrane and its trafficking, including endocytosis, is essential for autophagy (Imai *et al*, 2010; Noda *et al*, 2000; Puri *et al*, 2013; Young *et al*., 2006). Interestingly, genes involved in both autophagy and endocytosis have been associated with neurodegenerative disorders (Alegre-Abarrategui & Wade-Martins, 2009; Bandres-Ciga *et al*, 2019; Overhoff *et al*, 2020; Schreij *et al*, 2016; Vidyadhara *et al*, 2019). Loss of function of mammalian ATG9A *in vivo*, results in reduced autophagy, highlighting ATG9A’s importance in efficient autophagosome formation (Imai *et al*, 2016; Orsi *et al*., 2012; Popovic & Dikic, 2014; Saitoh *et al*, 2009). Brain-specific deletion of *Atg9a* in mice causes accumulation of the autophagy adaptor p62 and ubiquitinated proteins and swollen degenerative axons, linking loss of ATG9A to neurodegeneration (Yamaguchi *et al*, 2018).

Nuclear Dbf2-related (NDR) kinases NDR1 and NDR2 (NDR1/2) are highly-related AGC family serine/threonine kinases that are evolutionarily conserved from yeast to mammals (Hergovich *et al*, 2006). In various organisms they have been implicated in cellular proliferation (Leger *et al*, 2018), transcriptional asymmetry (Mazanka *et al*, 2008), DNA damage response (Qin *et al*, 2020), innate immunity (Liu *et al*, 2019), ciliogenesis (Chiba *et al*, 2013), neuronal differentiation (Emoto *et al*, 2004; Emoto *et al*, 2006; Rehberg *et al*, 2014; Ultanir *et al*, 2012), mitochondrial health (Wu *et al*, 2013) and autophagy (Amagai *et al*, 2015; Joffre *et al*, 2015; Martin *et al*, 2019). NDR kinases are required for dendritic arborisation and synapse formation in worms, flies and mice (Emoto *et al*., 2004; Gallegos & Bargmann, 2004; Ultanir *et al*., 2012) and substrates of NDR kinases, identified via chemical genetic methods, comprise of several membrane trafficking components (Ultanir *et al*., 2012). The NDR kinase orthologue wts is also required for the maintenance of dendrites in Drosophila (Emoto *et al*., 2006), yet the role of NDR kinases in neuronal health in mammals is not well-understood.

In order to investigate NDR1/2’s roles in neurons *in vivo*, we used *Ndr1* constitutive knockout and *Ndr2*-floxed mice (Schmitz-Rohmer *et al*, 2015) and knocked out *Ndr2* in excitatory neurons using the NEX-Cre driver (Goebbels *et al*, 2006). In dual neuronal *Ndr1* and *Ndr2* knockout mice, we observed prominent neurodegeneration in the cortex and hippocampus. Our comparison of hippocampal proteome and phosphoproteome between control and knockout littermates indicated a major alteration in endocytic pathways and revealed several novel NDR1/2 kinase substrates that contain the previously reported HXRXXS* motif. We validated Ras Association (RalGDS/AF-6) And Pleckstrin Homology Domains 1 (Raph1), a.k.a. Lamellipodin (Lpd) as a novel NDR1/2 substrate. We show that both NDR1/2 and Raph1 regulate neuronal endocytosis. In the absence of NDR1/2 p62 and ubiquitin accumulate in mouse neurons. Furthermore, autophagosome numbers are reduced in NDR1/2 knockout neurons and autophagic clearance is affected in primary neurons depleted of NDR1/2 kinases, overall showing that NDR1/2 are required for the maintenance of neuronal protein homeostasis. Finally, deletion of NDR1/2 in adult mice also results in a similar phenotype, indicating that NDR1/2 are needed for protein homeostasis and not only during neuronal development. Mechanistically, we showed that NDR kinases are critical regulators of clathrin-mediated endocytosis and ATG9A trafficking, revealing a novel kinase regulatory pathway that is essential for autophagy in neurons. We conclude that NDR1/2 are required for ATG9A trafficking, autophagy and for the maintenance of neuronal homeostasis to prevent neurodegeneration.

## Results

### Dual deletion of NDR1/2 kinases in excitatory neurons causes neurodegeneration and reduces mouse survival

To investigate the roles of NDR1/2 kinases in mammalian neurons *in vivo,* we generated a mouse model in which both *Ndr1* (also known as *stk38*) and *Ndr2* (also known as *stk38l*) were deleted in excitatory post-mitotic neurons from the cortex and hippocampus. Individual *Ndr1* or *Ndr2* full knockout mice are viable and fertile, but dual *Ndr1/Ndr2* knockout mice are embryonically lethal, indicating that the two isoforms compensate for each other’s function (Schmitz-Rohmer *et al*., 2015), likely owing to their 87% amino acid identity (Tamaskovic *et al*, 2003). Constitutive *Ndr1* knockout (*Ndr1*^KO^) and floxed *Ndr2* (*Ndr2*^flox^) mice (Schmitz-Rohmer *et al*., 2015) were crossed with mice expressing the Cre recombinase under the control of the NEX driver (Goebbels *et al*., 2006), which is specific for pyramidal neurons of the cortex and hippocampus. Our experimental crosses gave litters with 4 possible genotypes: *Ndr1*^KO/+^ *Ndr2*^flox/+^ NEX^Cre/+^ (control), *Ndr1*^KO/KO^ *Ndr2*^flox/+^ NEX^Cre/+^ (NDR1 KO), *Ndr1*^KO/+^ *Ndr2*^flox/flox^ NEX^Cre/+^ (NDR2 KO) and *Ndr1*^KO/KO^ *Ndr2*^flox/flox^ NEX^Cre/+^ (NDR1/2 KO). As previously reported (Cornils *et al*, 2010), NDR1 KO mice were viable, fertile and exhibited normal brain development and the same was true of the NDR2 KO mice, which lacked NDR2 in excitatory neurons (Fig. S1A &S1B). NDR1/2 KO mice were also viable but had significantly lower weights and reduced survival rate compared to littermates (Fig. 1A &1B). Weight and survival rate were not affected in NDR1 or NDR2 individual KO mice (Fig. 1A &1B), further confirming their mutual compensation.

**Figure 1.**
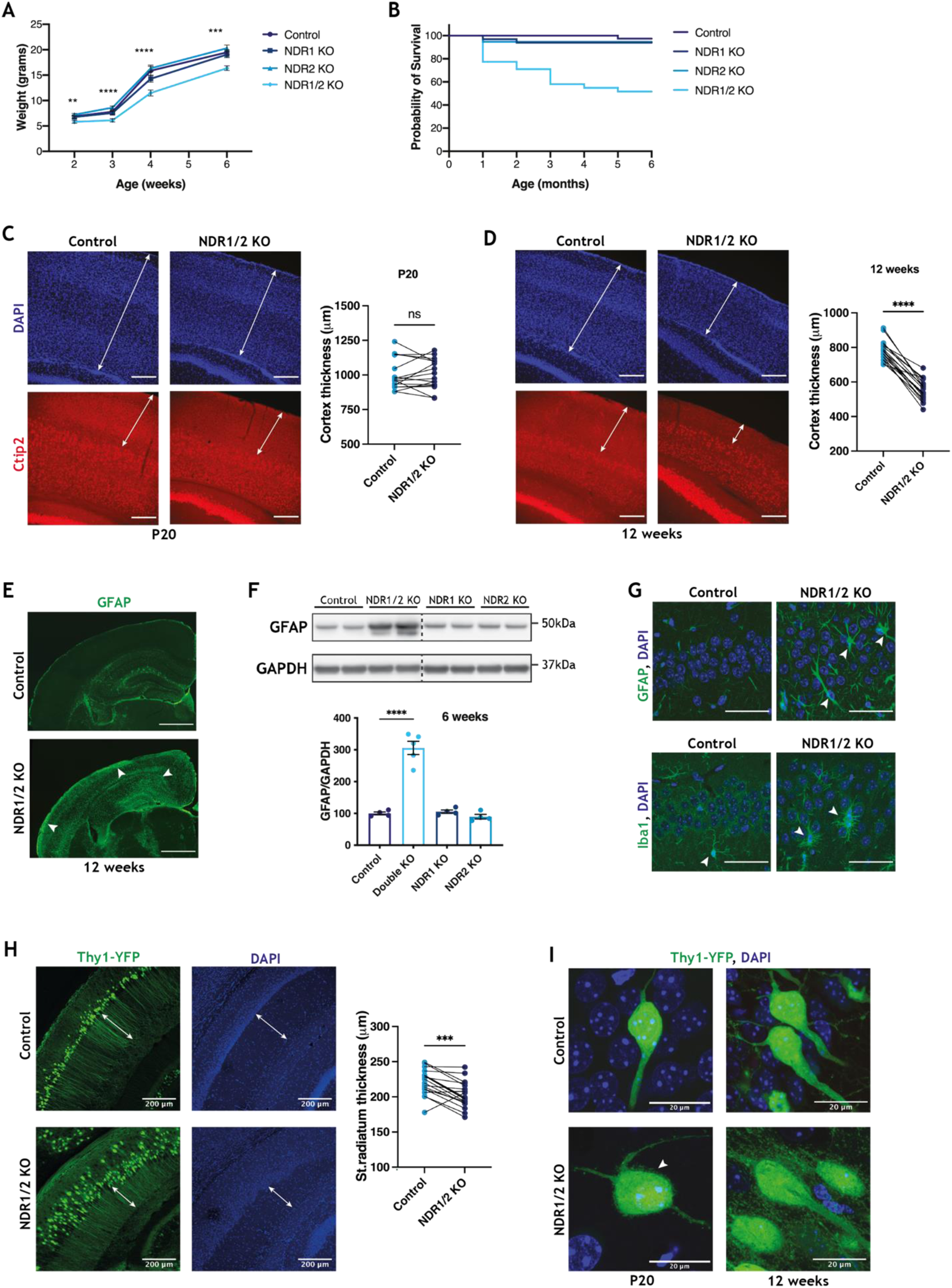
Dual loss of NDR kinases in neurons leads to neurodegeneration. **(A)** Line graph showing the average weight of NDR1 KO, NDR2 KO, NDR1/2 KO and control mice up to 6 weeks of age. The differences in weight between the genotype groups at each time-point were analysed using ordinary one-way ANOVAs or Kruskal-Wallis tests, n = 20-30 mice/ group. **(B)** Graph illustrating the probability of survival of NDR1 KO, NDR2 KO, NDR1/2 KO and control mice up to 6 months of age. n = 20-30 mice/ group. **(C, D)** Immunofluorescence staining of Ctip2 in brain slices of NDR1/2 KO and control mice at P20 or 12 weeks of age. White arrowed lines in the DAPI images show the thickness of the cortex and white arrowed lines in the Ctip2 images show the thickness of the upper layers I-IV of the cortex. Scale bars 200μm. The graphs show quantifications of cortex thickness and the data was analysed using paired Student’s *t* tests, n = 18 measurements from 3 mice/ genotype. **(E)** Immunofluorescence staining of GFAP in brain slices of 12-week-old NDR1/2 KO and control mice. White arrows show areas with increased GFAP signal in NDR1/2 KO mice. **(F)** Western blot analyses of GFAP levels in lysates from the cortex of 6-week-old mice. GAPDH was used as a loading control. The graphs show quantifications of the GFAP bands normalised against the GAPDH levels and the data was analysed using an ordinary one-way ANOVA with Tukey’s post hoc test, n = 3-5 mice/ group. **(G)** Immunofluorescence staining of GFAP and the microglial marker Iba1 in the CA1 area of the hippocampus. White arrows indicate cells expressing the above-mentioned markers. Scale bars 50μm. **(H**) Images from brain slices of 12-week-old Thy1-YFP expressing mice in the CA1 area of the hippocampus. White arrowed lines indicate the stratum radiatum, where CA1 neuron dendrites are visible in YFP. The graph shows quantification of stratum radiatum thickness and the data was analysed using a paired Student’s *t* test, n = 18 measurements from 3 mice/ genotype. **(I)** Images from brain slices of Thy1-YFP expressing mice in the CA1 cell body layer. The white arrow shows membrane protrusions present in NDR1/2 knock-out neurons.

In NDR1/2 KO mice cortical thickness and layering were unchanged at postnatal day 20 (P20) (Fig. 1C), but at 12 weeks of age the cortex thickness was significantly reduced (Fig. 1D). Immunostainings of the cortical layer V-VI nuclear marker Ctip2 showed that layers V-VI were still present in 12-week-old NDR1/2 KO mice, indicating a specific loss of the upper layers I-IV (Fig. 1D). Individual NDR1 or NDR2 KOs were not affected, even as late as 20 weeks of age (Fig. S1A). Astrocyte and microglial activation are hallmarks of neurodegeneration in humans and mouse models. The astrocytic marker GFAP was highly increased in NDR1/2 KO mice (Fig. 1E, 1F & S1C) and additionally, microglia exhibited a hypertrophied morphology associated with ‘reactive’ cells, in contrast to the ramified ‘resting’ microglial cells (Ransohoff, 2016) present in control mice (Fig. 1G). Individual NDR1 KO and NDR2 KO mice did not have increased GFAP levels, despite a clear reduction in the protein levels of NDR1 and NDR2, respectively (Fig. 1F, S1B & S1C). H&E stainings from the upper layers of the cortex of NDR1/2 KOs revealed neurons with a deeply eosinophilic cytoplasm and small condensed nuclei, indicative of apoptosis or necrosis (Fig. S1D). To a lesser extent, necrotic neurons were also observed in the hippocampus (Fig. S1D).

To explore the morphology of these degenerating neurons, we crossed NDR1/2 KO mice with Thy-YFP mice, which express YFP sparsely in pyramidal neurons (Feng *et al*, 2000). In the hippocampus, the CA1 cell body layer was disorganised and stratum radiatum thickness was significantly reduced, indicating neuropil loss (Fig. 1H). Overall, our data shows that dual, but not individual loss of NDR1 and NDR2 in neurons leads to loss of upper cortical layers and neurodegeneration of the cortex and hippocampus.

NDR1/2 play a role in dendrite and spine morphogenesis and actin regulation (Emoto *et al*., 2004; Geng *et al*, 2000; Ultanir *et al*., 2012). NDR1/2 knockout neurons have small protrusions and membrane ruffles on the cell body and dendrites, in contrast to the smooth plasma membrane of control cells at postnatal day 20 (Fig. 1I). By 12 weeks these protrusions become more exuberant and occupy the area in between the cells, as the knockout neurons lose the tightly packed distribution of cell bodies characteristic of control neurons (Fig. 1H&1I). Despite this, we observed no differences in the total levels of key synaptic proteins in NDR1/2 KO mice (Fig. S1E). Transmission electron microscopy (TEM) assessment of brain slices from NDR1/2 KO mice confirmed the changes in cellular morphology seen in Thy1-YFP mice and revealed changes in mitochondria, which were rounder and more fragmented, compared to control brains (Fig. S1F). Such changes in mitochondrial morphology have been previously reported in apoptotic cells and neurodegenerative conditions (Knott *et al*, 2008; Su *et al*, 2010), and could represent an early step during the apoptosis process, but could also be a result of the more direct role of NDR kinases in mitochondrial quality control (Wu *et al*., 2013). In this study we aimed to uncover the mechanisms through which loss of both NDR1/2 kinases impinges on neuronal health in mice.

### Proteomics analyses of NDR1/2 KO hippocampi show changes in endocytosis-related proteins and reveal novel substrates

To gain mechanistic insight into the function of NDR1/2, we employed a mass spectrometry approach to identify the substrates of NDR that could mediate their roles and to assess changes in the levels of proteins in NDR1/2 KO brains at a global scale. Due to the high GFAP expression in the cortex of NDR1/2 KO mice as early as P20 (Fig. S1C), we used hippocampus samples for the proteomics analysis. Astrocytes are not targeted by the Cre recombinase and could confound our results, since we are interested in neuron-specific changes that happen directly downstream of NDR1/2 loss. Briefly, hippocampi were dissected from NDR1/2 KO and control littermates. A total of 5 control and 5 NDR1/2 KO samples were trypsin digested and labelled with a TMT (Tandem Mass Tag) 10-plex reagent (Jiang *et al*, 2017). For phospho-proteome analysis, phospho-peptides were enriched using the titanium dioxide and Fe-NTA methods (Jiang *et al*., 2017) (Fig. 2A).

**Figure 2.**
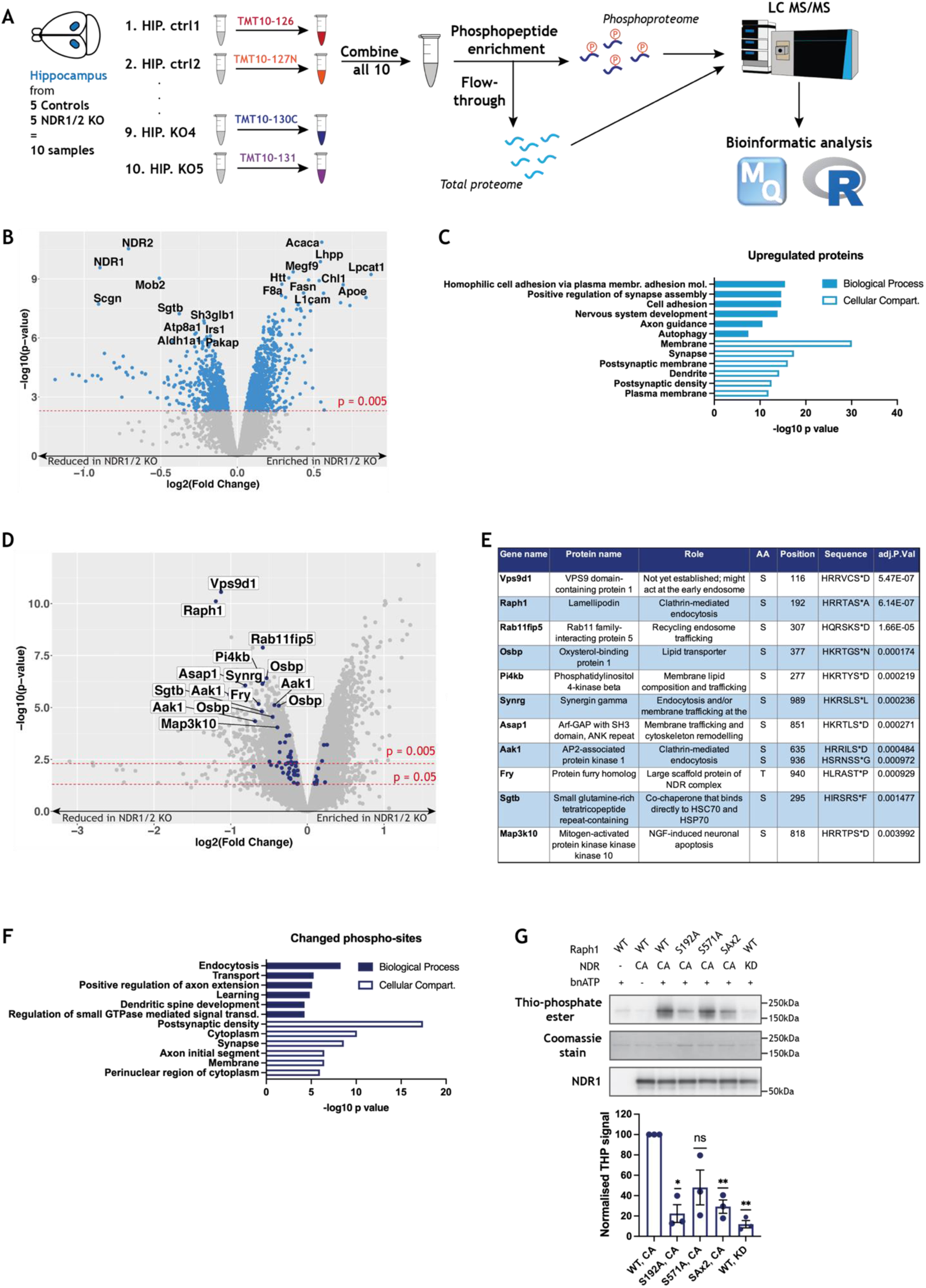
Quantitative proteomics and phospho-proteomics indicate altered endocytosis and identify Raph1/ Lpd as an NDR kinase substrate. **(A)** Workflow of Tandem Mass Tag (TMT) labelling of hippocampus samples from the brains of NDR1/2 knock-out and control mice with subsequent phospho-enrichment and mass spectrometry analysis of the total and the phospho-proteome. Max Quant software and R coding were used for the bioinformatic analysis. **(B)** Volcano plot of the difference in protein levels between control and NDR1/2 knock-out mice. Each point represents one protein. The x axis shows log2 transformed fold change and the y axis shows significance by −log10 transformed p-value obtained by linear models for microarray data (LIMMA). All significantly changed proteins are highlighted in blue. A protein is considered significantly changed if its p-value is <0.005, with the red line marking the difference between significantly and non-significantly changed proteins. The top 10 up-regulated or down-regulated proteins in the NDR1/2 knock-out brains compared to the control brains are labelled in black. **(C, E)** Results from gene enrichment analyses run using the online tool Database for Annotation, Visualization and Integrated Discovery (DAVID), with the list of genes corresponding to all significantly upregulated proteins (C) or all significantly changed phospho-peptides (E). The x axis shows the p-value or EASE score generated by DAVID to show how enriched a term associated with a list of genes is. The top 6 most enriched terms from each category are represented. (D) Volcano plot showing the difference in phospho-peptide levels between control and NDR1/2 knock-out mice. Each point represents one phospho-peptide. The x axis shows log2 transformed fold change and the y axis shows significance by −log10 transformed p-value obtained by LIMMA. All phospho-peptides above the higher red line have a p-value <0.005 and all phospho-peptides above the lower red line have a p-value <0.05. The blue dots represent phospho-peptides with the already established NDR consensus motif HXRXXS/T. All phospho-peptides with the NDR consensus motif and an adjusted p-value <0.005 have been labelled. **(F)** Table with the roles, specific phosphorylation site and phosphorylated sequence of all NDR substrate candidates with an adjusted p-value <0.005. **(G)** Representative western blots showing thio-phosphate ester and NDR1 levels in an *in vitro* kinase assay. Total Raph1 levels are shown by Coomassie staining. The bar graph shows quantification of Raph1 thio-phosphorylation normalised to total Raph1 levels and expressed as a percentage of the Raph1-WT/ NDR-CA thio-phosphate ester level. The data was analysed using a one sample *t* and Wilcoxon. n=3 independent experiments.

The total proteome analysis identified 7456 proteins, out of which 408 were significantly upregulated and 255 were downregulated in the NDR1/2 KO brains (Fig. 2B & Supplemental Data 1). Remarkably, the top two most reduced proteins were NDR1 and NDR2, as expected (Fig. 2B). Interestingly, Mob2, a conserved interactor and activator of NDR kinases (Bichsel *et al*, 2004; Weiss *et al*, 2002) was also markedly downregulated in the absence of its interacting partners NDR1/2. The top upregulated proteins include several enzymes involved in lipid metabolism, such as Lpcat, Acaca and Fasn and the cell adhesion molecules L1cam and Chl1 (Fig. 2B). Accordingly, when running gene enrichment analyses using the online tool DAVID (Huang da *et al*, 2009a, b), ‘cell adhesion’ comes up a biological process highly enriched in the list upregulated proteins. In addition, ‘membrane’-related terms are also enriched in the cellular compartment category, likely matching the changes in lipid metabolism (Fig. 2C & Supplemental Data 2). Another biological process associated with the list of upregulated proteins that stands out is “autophagy”, as it is already established that NDR1/2 play a role in autophagy in mammalian cells (Amagai *et al*., 2015; Joffre *et al*., 2015; Martin *et al*., 2019) (Fig. 2C & Supplemental Data 2). The significantly downregulated proteins were associated with biological processes and cellular compartments linked to myelination (Fig. S2A & Supplemental Data 3). Interestingly, a reduction in myelination can be indicative of axonal degeneration, which has been previously reported in mouse models with neuron-specific deletions of autophagy-related proteins (Komatsu *et al*, 2007b; Liang *et al*, 2010; Zhao *et al*, 2013). In terms of KEGG pathways and human phenotype ontology, the enriched terms within these categories describe features or diseases reminiscent of the neurodegeneration phenotype observed in NDR1/2 KO brains, such as “neurofibrillary tangles”, “dementia” and “cerebral inclusion bodies” (Fig. S2B & Supplemental Data 4), as would be expected. Given the established role of NDR1 in autophagy (Joffre *et al*., 2015) and the role of autophagy in neurodegeneration (Hara *et al*., 2006; Komatsu *et al*., 2006), these analyses support that NDR1/2 could be involved in neuronal autophagy.

To identify signalling differences between control and NDR1/2 KO mice, we compared their hippocampal phosphoproteomes. TMT labelling mass spectrometry identified 39620 unique phosphorylation sites, with numerous significant differences between the two genotypes (Fig. 2D). We used the established NDR kinase consensus motif HXRXXS/T (Mazanka *et al*., 2008; Ultanir *et al*., 2012) to filter the data for potential NDR substrates and applied a stringent significance cut-off of adjusted p value <0.005 to find the most robustly reduced phospho-sites in the absence of NDR1/2 kinases (Fig. 2D & Supplemental Data 5). This produced a list of 11 putative NDR1/2 substrates (Fig. 2D &2E). Three of them were previously reported to be direct substrates of NDR1/2 in brains, namely PI4KB, Rab11fip5 and AAK1 (Ultanir *et al*., 2012), providing validation to our experiment. Most of the substrates play roles in membrane trafficking (Fig. 2E). To assess the implications of all changed phosphorylations, we performed gene enrichment analyses using the genes corresponding to altered phosphorylation sites with a p value <0.005 (Fig. 2F). Interestingly, ‘endocytosis’ was the most significant biological process enriched in this list of genes, providing a link between NDR1/2 kinases and this cellular process (Fig. 2F & Supplemental Data 6). Furthermore, the putative direct substrates Raph1, AAK1 and SYNG all regulate clathrin-mediated endocytosis (CME) (Conner & Schmid, 2002; Hirst *et al*, 2005; Vehlow *et al*, 2013). Collectively, phosphoproteomics analysis strongly indicate that NDR1/2 kinases play a role in membrane trafficking and endocytosis (Fig. 2E).

We decided to validate one of the most promising substrate candidates, Raph1/Lamellipodin (Lpd), a protein that interacts with endophilin A and plays roles in clathrin-mediated endocytosis and fast endophilin-mediated endocytosis (Boucrot *et al*, 2015; Chan Wah Hak *et al*, 2018; Vehlow *et al*., 2013). Interestingly, like NDR kinases Lpd is also required for neuronal dendritic arborisation (Tasaka *et al*, 2012). Lpd harbors a Ras association (RA) domain followed by a Pleckstrin homology (PH) domain (Fig. S2C) (Krause *et al*, 2004). The phospho-site identified with mass spectrometry, Ser192, precedes the RA domain and we noted that another residue with the NDR consensus motif, Ser571, is located closely after the PH domain (Fig. S2C). We purified single and double phosphomutant Lpd (S192A & S571A) (Fig. S2D) and performed *in vitro* kinase assays using constitutively active (CA) NDR1 or kinase dead (KD) NDR1 with wild-type or phosphomutant Raph1/Lpd (Fig. 2G &S2E). We found that NDR1 specifically phosphorylates Raph1/Lpd on S192, while S571 is not consistently phosphorylated *in vitro* (Fig. 2G). Phosphorylation of S192 was confirmed in HEK293T cells with a phospho-specific antibody raised against a phospho-peptide containing this site (Fig. S2F). Overall, our phosphoproteomics results indicate that NDR1/2 play a role in endocytosis and identify Raph1/Lpd as a novel substrate.

### NDR1/2 depletion reduces endocytosis, partially through Raph1/ Lpd

Due to the significant changes in endocytosis-related phosphorylation events in our proteomics analysis, we wanted to assess whether NDR1/2 depletion impacts endocytosis in cultured primary neurons. Transferrin receptor (TfR) is a transmembrane protein that internalises transferrin (Tf)-bound iron and subsequently gets recycled through the endosomal pathway back to the plasma membrane to ensure further iron uptake (Aisen, 2004). Transferrin-based assays are used to study clathrin mediated endocytosis. We performed labelled Tf-uptake experiments, using transferrin conjugated to Alexa-Fluor 568 (Tf-568) in hippocampal neurons previously infected with either scramble shRNA or dual NDR1 shRNA and NDR2 shRNA, which robustly knocked-down their target genes (Fig. S3A). After 20 min of incubation with Tf-568 (pulse), the Tf-containing media was replaced with maintenance media devoid of Tf-568, for 20 min or 60 min (chase), to allow for recycling of transferrin. Tf-568 signal was significantly reduced in NDR1/2 shRNA neurons after pulse, indicating that endocytosis is less efficient in the absence of NDR1/2 kinases (Fig. 3A). No significant difference was detected between scramble shRNA and NDR1/2-depleted neurons after the 20 min Tf chase, indicating that by this point scramble shRNA cells had recycled more Tf, since they had taken up significantly more Tf to begin with (Fig. 3A).

**Figure 3.**
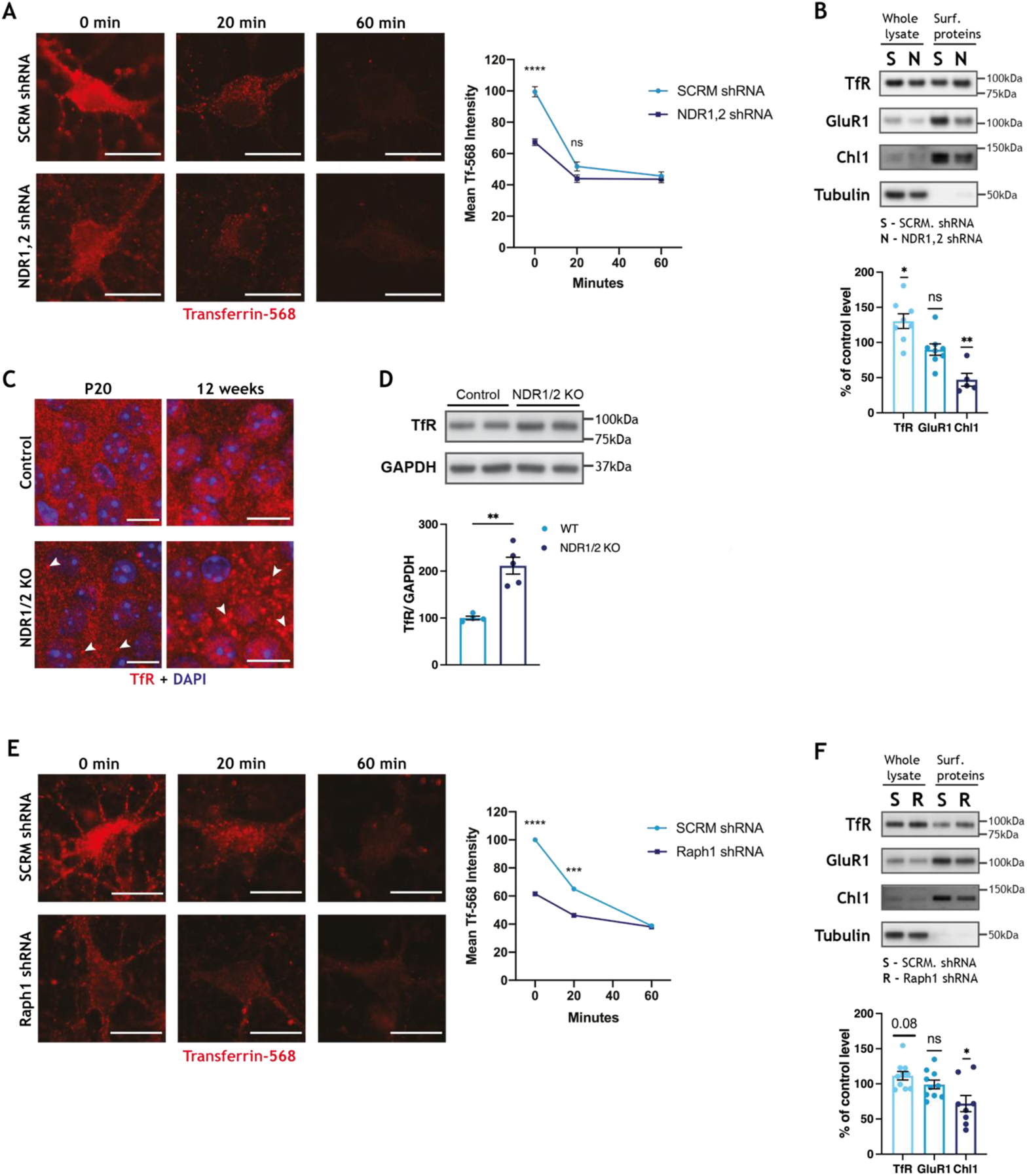
NDR kinases and their substrate Raph1/Lpd are required for endocytosis. **(A & E)** Representative images of DIV11 rat hippocampal primary neurons incubated with transferrin-Alexa568 (transferrin-568) for 20 min (pulse) and chased for 0, 20, or 60 min in complete media. Scale bars 10μm. Neurons were infected with scramble (SCRM) shRNA lentivirus or lentiviral vectors expressing NDR1 and NDR2 shRNAs (NDR1,2 shRNA) (A) or Raph1/Lpd shRNA (E) and transfected with an empty EGFP-expressing plasmid (not depicted). Line graphs represent the quantification of cellular transferrin-568 normalised to cell area (using the EGFP cell fill) after the chase period. The data was analysed using a mixed-effects analysis with Šidák’s multiple comparisons test. n>30 cells/ condition from 3 independent experiments. **(B & F)** Western blot analyses of surface biotinylation experiments on DIV12 rat cortical neurons infected with scramble SCRM or NDR1,2 shRNA lentivirus (B) or Raph1 shRNA (F). Surface protein levels were normalised against input. Bar graphs show surface protein levels expressed as a percentage of the corresponding SCRM shRNA control level. The data was analysed using a one sample *t* and Wilcoxon test. n=6 samples/ condition from 3 independent experiments. **(C)** Immunofluorescence staining of transferrin receptor (TfR) in the CA1 area of the hippocampus in brain slices from P20 and 12-week-old NDR1/2 knock-out and control mice. White arrows indicate TfR-positive puncta. Scale bars 10μm. **(D)** Western blot analyses of TfR in lysates from the cortex of 6-week-old NDR1/2 knock-out and control mice. GAPDH was used as a loading control. The bar graph shows quantifications of the TfR bands normalised against the GAPDH levels and the data analysed using an unpaired Student’s *t* test, n = 4-5 mice/ group.

To confirm an impairment in TfR trafficking, we assessed the levels of TfR at the plasma membrane using surface biotinylation experiments. Biotin labelling of surface proteins was carried out on ice to block endocytosis and biotinylated proteins were subsequently isolated and their levels measured using western blotting. The surface levels of TfR were higher in neurons in which NDR1/2 had been knocked down when normalised to total TfR levels (Fig. 3B). Surprisingly, the surface levels of the adhesion molecule Chl1, a regulator of integrin signalling (Buhusi *et al*, 2003) which was significantly upregulated in the brains of NDR1/2 KO mice (Fig. 2B), was reduced in NDR1/2-depleted neurons (Fig. 3B). The NDR kinase homolog trc is required for adhesion of dendrites to epithelia via integrin signalling in flies (Han *et al*, 2012), so it is possible that mammalian NDR1/2 has a similar function, mediating neuronal morphology and cell body layering in mammals. By contrast, surface levels of the AMPA-type glutamate receptor subunit GluR1 were not altered due to NDR1/2 depletion (Fig. 3B), in agreement with no apparent differences in the levels of synaptic markers in NDR1/2 KO brains (Fig. S1E). These results suggest that both TfR and Chl1 trafficking, and consequently their surface levels, are impaired in the absence of NDR1/2.

To check if TfR trafficking is altered *in vivo* in NDR1/2 KO mice, we assessed the localisation of endogenous TfR in mouse brains using immunohistochemistry. Compared to the diffuse and less discrete TfR signal in control mice, NDR1/2 KO brains contained distinct TfR puncta, that accumulated with age, indicative of a blockage in TfR recycling (Fig. 3C). At P20 few TfR puncta were present in NDR1/2 KOs, but these increased to a striking number by 12 weeks of age (Fig. 3C), making TfR accumulation an early impairment that increases progressively. The increase in TfR was confirmed with western blots at 6 weeks of age (Fig. 3D). To characterise TfR puncta in NDR1/2 KOs, we co-stained TfR with endosomal markers. The retromer component VPS35 (Seaman, 2012), which plays a role in tubulation of endosomes, also displayed a similar discrete accumulation and robustly colocalised with TfR puncta in NDR1/2 KOs (Fig. S3B). Taken together, these results show that there is an accumulation of TfR and VPS35 positive endosomal compartments in NDR1/2 KO neurons, highlighting defects in endosomal trafficking.

Our phosphoproteome assessment identified several NDR1/2 target substrates which could contribute to the membrane trafficking defects highlighted above. We selected Raph1, a known mediator of endocytosis (Vehlow *et al*., 2013), and tested if it plays a role in TfR trafficking, using lentivirus to knockdown Raph1 in primary neurons (Fig. S3C). Endocytosis of Tf-568 was significantly reduced in Raph1 knock-down neurons compared to the scramble shRNA control, mimicking loss of NDR1/2 (Fig. 3E). After a 20 min chase the Raph1shRNA neurons had recycled less Tf, but this was proportional to their lower Tf uptake after pulse, indicating that Raph1 likely does not impact Tf recycling (Fig. 3E). Interestingly, although the surface levels of TfR were also increased in Raph1 shRNA neurons, this change was not significant (Fig. 3F), indicating that NDR may be acting via multiple substrates to regulate TfR trafficking. Chl1 surface levels were significantly reduced, while GluR1 was not affected in the absence of Raph1 (Fig. 3F), mimicking NDR1/2 knockdown. Overall, these results suggest that NDR1/2 and their substrate Raph1 regulate endocytosis of TfR and alter membrane trafficking.

### Autophagy is impaired in neurons that lack NDR1/2 kinases

Our NDR1/2 KO mouse model presented striking similarities to brain specific autophagy defective mouse models, such as the timing of neuronal loss and specific loss of upper cortical layers (Hara *et al*., 2006; Komatsu *et al*., 2006). Furthermore, several autophagy-related proteins are upregulated in our proteomics dataset and changes in endocytosis can impact autophagosome formation (Popovic & Dikic, 2014; Puri *et al*., 2013; Tooze *et al*, 2014). Given the reported role of NDR1/2 kinases in autophagy, we decided to test if autophagy is impaired in our mouse model. The autophagy adaptor p62 binds ubiquitinated proteins and targets them to autophagosomes, while also being itself degraded by autophagy (Komatsu *et al*, 2007a; Pankiv *et al*, 2007). Immunostainings in brain sections revealed a prominent accumulation of p62 in NDR1/2 KOs by 12 weeks of age (Fig. 4A). On western blots it became obvious that at P20 the increase in p62 is not yet significant, but by 6 weeks of age significantly more p62 is present in NDR1/2 KO brains (Fig. 4B), indicating that there is a gradual age-dependent increase in this marker.

**Figure 4.**
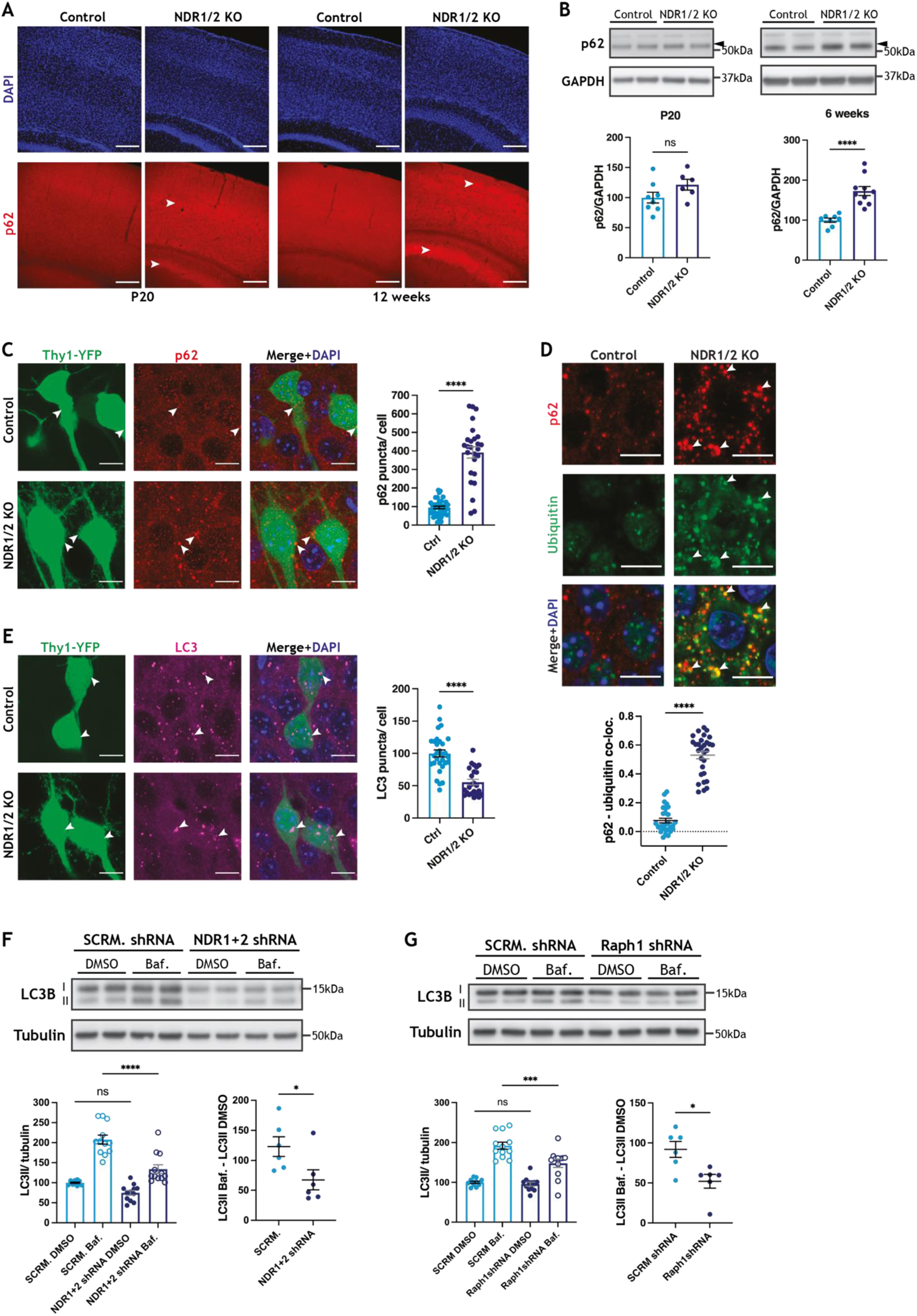
Autophagy is impaired in NDR1/2 KO mice. **(A)** Immunofluorescence staining of the autophagy receptor p62 in brain slices of NDR1/2 KO and control mice at P20 and 12 weeks of age. White arrows indicate areas of increased p62 signal. Scale bars 200μm. **(B)** Western blot analyses of p62 levels in lysates from the cortex of P20 and 6-week-old NDR1/2 KO and control mice. GAPDH is used as a loading control. The bar graphs show quantifications of the p62 bands normalised against the GAPDH levels and the data was analysed using unpaired Student’s *t* tests, n = 3-5 mice/ group, 2 technical replicates. **(C, E)** Immunofluorescence staining of p62 (C) and LC3 (E) in the CA1 area of the hippocampus in brain slices from 12-week-old Thy1-YFP expressing mice. Scale bars 50μm. The bar graphs show the number of p62 (C) or LC3 (E) puncta in neurons normalised against the cell body area using the YFP signal. The data was analysed using a Mann-Whitney test for p62 and an unpaired Student’s *t* test for LC3. n = 38 control and 27 NDR1/2 KO neurons from 2 mice per genotype for p62. n=29 control and 24 NDR1/2 KO neurons from n=2 mice per genotype for LC3. **(D)** Immunofluorescence staining of p62 and ubiquitin in CA1 at 12 weeks. White arrows indicate co-localisation between p62 and ubiquitin puncta. Scale bars 10μm. The scatter plot shows quantification of p62 and ubiquitin co-localisation expressed as Pearson correlation coefficient. The data was analysed with a Mann Whitney test. n=30 measurements from 3 mice/ genotype. **(F-G)** Western blot analyses of LC3 levels in lysates from DIV13 rat primary cortical neurons infected with lentiviruses expressing a scramble (SCRM) shRNA and shRNAs targeting NDR1 and NDR2 (F) or Raph1(G). The cells were treated with DMSO or 100nM of Bafilomycin A1 (Baf.) for 4 hours prior to lysis. Tubulin was used as a loading control. The bar graphs show quantification of the LC3II bands normalised against the tubulin levels and the data was analysed using ordinary one-way ANOVAs with Tukey’s post hoc test. n = 6 samples/ group from 3 independent experiments, 2 technical replicates. The scatter plots show quantifications of the absolute increase in LC3II between the DMSO and the Bafilomycin A1 conditions. n = 6 measurements/ group from 3 independent experiments, 2 technical replicates.

Considering the significant astrogliosis in NDR1/2 KO brains, we wanted to establish if the observed accumulation in p62 was specifically present in neurons or in non-neuronal cells. To this end we used Thy1-YFP expressing mice and quantified p62 puncta in YFP-positive cell bodies. The results revealed a significant increase in p62 puncta in the soma of NDR1/2 KO neurons at 12 weeks of age (Fig. 4C). Furthermore, stainings from brain sections also showed a marked increase in cytoplasmic ubiquitin in NDR1/2 KO neurons and ubiquitin puncta colocalised with p62, revealing an accumulation in ubiquitinated proteins targeted to the autophagy pathway (Fig. 4D). The increase in ubiquitinated proteins was also confirmed with western blots in the brains of 6-week-old NDR1/2 KO mice (Fig. S4A). These results agree with increased ubiquitin and p62 accumulations in autophagy-impaired mouse models (Hara *et al*., 2006; Komatsu *et al*., 2006; Kuijpers *et al*, 2021).

The accumulation in p62 and ubiquitinated proteins could result from a defect in autophagosome formation and subsequent impairment in the clearance of these proteins. Consequently, we assessed whether the levels of the autophagosome marker LC3 were changed in NDR1/2 KO neurons. We detected autophagosomes in YFP-positive neuronal cell bodies and found that the number of LC3 puncta was significantly lower in NDR1/2 KO neurons when compared to controls (Fig. 4E). These results indicate that autophagosome formation is reduced in the absence of NDR kinases.

If autophagosomes still form, but ubiquitinated proteins are not cleared adequately, we wondered if this could be a result of inefficient targeting of ubiquitinated proteins to the autophagy pathway or a failure of autophagosomes to fuse with lysosomes. We performed co-stainings of p62, LC3 and the lysosomal marker LAMP2 in brain sections. In both controls and NDR1/2 KOs almost all LC3 puncta colocalised with p62 and LAMP2 (Fig. S4B), indicating that LC3 staining represents autolysosomes in the soma. Larger cytoplasmic p62 puncta colocalised with LC3 and LAMP2 in controls, showing targeting of p62 to these compartments (Fig. S4B). By contrast, while some p62 colocalised with LC3 and LAMP2 in NDR1/2 KO neurons similarly to controls (Fig. S4B - white arrows), there was a substantial amount of p62 puncta that did not colocalise with either LC3 or lysosomes (Fig. S4B - yellow dotted circles). Considering the gradual accumulation of p62 we observed at P20, 6 weeks and 12 weeks of age, we suggest that autophagy functions at reduced levels, which results in accumulation of ubiquitinated proteins in NDR1/2 KO neurons over a longer period.

To test this hypothesis, we treated primary neurons with Bafilomycin A1, a drug that inhibits autophagosome-lysosome fusion, blocking autophagosome clearance and enabling assessment of formation rates (Mizushima & Yoshimori, 2007; Rubinsztein *et al*, 2009). We assessed autophagosome formation in rat primary cortical neurons infected with lentiviral vectors expressing NDR1 and NDR2 shRNAs or a scramble shRNA control. Bafilomycin A1 treatment resulted in a significant increase in lipidated LC3 (LC3II) levels in both scramble shRNA control and NDR1/2-depleted neurons (Fig. 4F), indicating that autophagosomes formed in both conditions. However, LC3II levels after Bafilomycin A1 treatment were significantly lower in the NDR1/2 shRNA-infected neurons and the amount of LC3II that accumulated within the treatment time (LC3II Baf. – LC3II DMSO) was also reduced (Fig. 4F). These results show that both the formation and degradation of autophagosomes are reduced in NDR1/2-depleted neurons. LC3I levels were also reduced in NDR1/2 KO neurons (Fig. S4C), and this may be due to indirect mechanisms downstream of NDR1/2, which impinge on LC3I synthesis. We next assessed if the role of NDR was dependent on its kinase activity. For this, we transfected HEK293T cells with either constitutively active NDR1 (NDR1-CA) or a kinase-dead version of NDR1 (NDR1-KD) (Ultanir *et al*., 2012) and treated them with Bafilomycin A1. While basal levels of LC3II were not changed in the NDR1-CA or NDR1-KD expressing cells, the levels of LC3II post-Bafilomycin treatment were significantly higher with NDR1-CA and reduced with NDR1-KD (Fig. S4D). These results suggest that the kinase activity of NDR1 is necessary for efficient autophagy.

Consequently, we tested if knockdown of the newly validated NDR substrate Raph1 could mimic some of the effects seen with NDR1/2 knockdown on autophagosome levels. Knockdown of Raph1 in neurons also resulted in a reduction in LC3II accumulation upon Bafilomycin A1 treatment (Fig. 4G), without any changes in LC3I (Fig. S4E), indicating that Raph1 could act downstream of NDR1/2 to mediate its role in autophagy. These results show that NDR1 and NDR2, are required for efficient autophagy in neurons, and via their kinase activity, NDR kinases are sufficient to enhance autophagy in mammalian cells.

### ATG9A trafficking and localisation is altered in NDR1/2 KO mice

TfR-positive recycling endosomes have been linked to the autophagy pathway (Lamb *et al*, 2013; Puri *et al*., 2013; Soreng *et al*, 2018; Tooze *et al*., 2014). Early-acting key autophagy proteins like ATG9A and ULK1 can be found on TfR-positive recycling endosomes and transferrin-containing recycling endosomal membranes can contribute to the formation of new autophagosomes (Longatti *et al*, 2012). In mammalian cell lines ATG9A can be present at the plasma membrane (Puri *et al*., 2013; Zhou *et al*, 2017), where it gets internalised via CME and delivered to the early endosome. From here it is sorted to the recycling endosome, before reaching autophagosome initiation sites, to contribute membrane to autophagosome precursors (Puri *et al*., 2013). A block in endocytosis impairs autophagy and results in ATG9A redistribution from perinuclear areas to areas close to the plasma membrane, concomitant with a reduction in ATG9A colocalisation with either Golgi or recycling endosome markers (Puri *et al*., 2013). Due to severe alterations in TfR and endosomes, we hypothesised that the function of the transmembrane protein ATG9A may be impaired, contributing to the observed autophagy deficits. Under normal conditions ATG9A cycles between membrane compartments, such as the Golgi, recycling endosomes and the plasma membrane and upon autophagy induction it relocates to peripheral sites of autophagosome formation; ATG9A-positive vesicles provide membrane to the growing phagophore, which will eventually close to form the autophagosome (Judith *et al*., 2019; Longatti *et al*., 2012; Orsi *et al*., 2012; Soreng *et al*., 2018; Young *et al*., 2006). In control brains, ATG9A had a perinuclear localisation, as expected, while in NDR1/2 KOs ATG9A localised to more distal dendritic regions and had a more punctate distribution (Fig. 5A). Interestingly, the levels of ATG9A were also increased in the brains of NDR1/2 KO mice at 12 weeks of age (Fig. 5B), even though at P20 ATG9A levels were unchanged (Supplemental Data 1), indicating that there is a chronic accumulation of this autophagy component. We co-stained ATG9A with a well-established Golgi marker, GM130, and found that while in control neurons the vast majority of ATG9A colocalises with GM130, colocalisation is significantly reduced in NDR1/2 KO neurons in the cell body layer of CA1 (Fig. 5C). Interestingly, ATG9A mislocalisation was observed as early as P20 (Fig. S5A), supporting the idea that defects in ATG9A and TfR trafficking, downstream of NDR substrates, precede accumulation of p62 aggregates.

**Figure 5.**
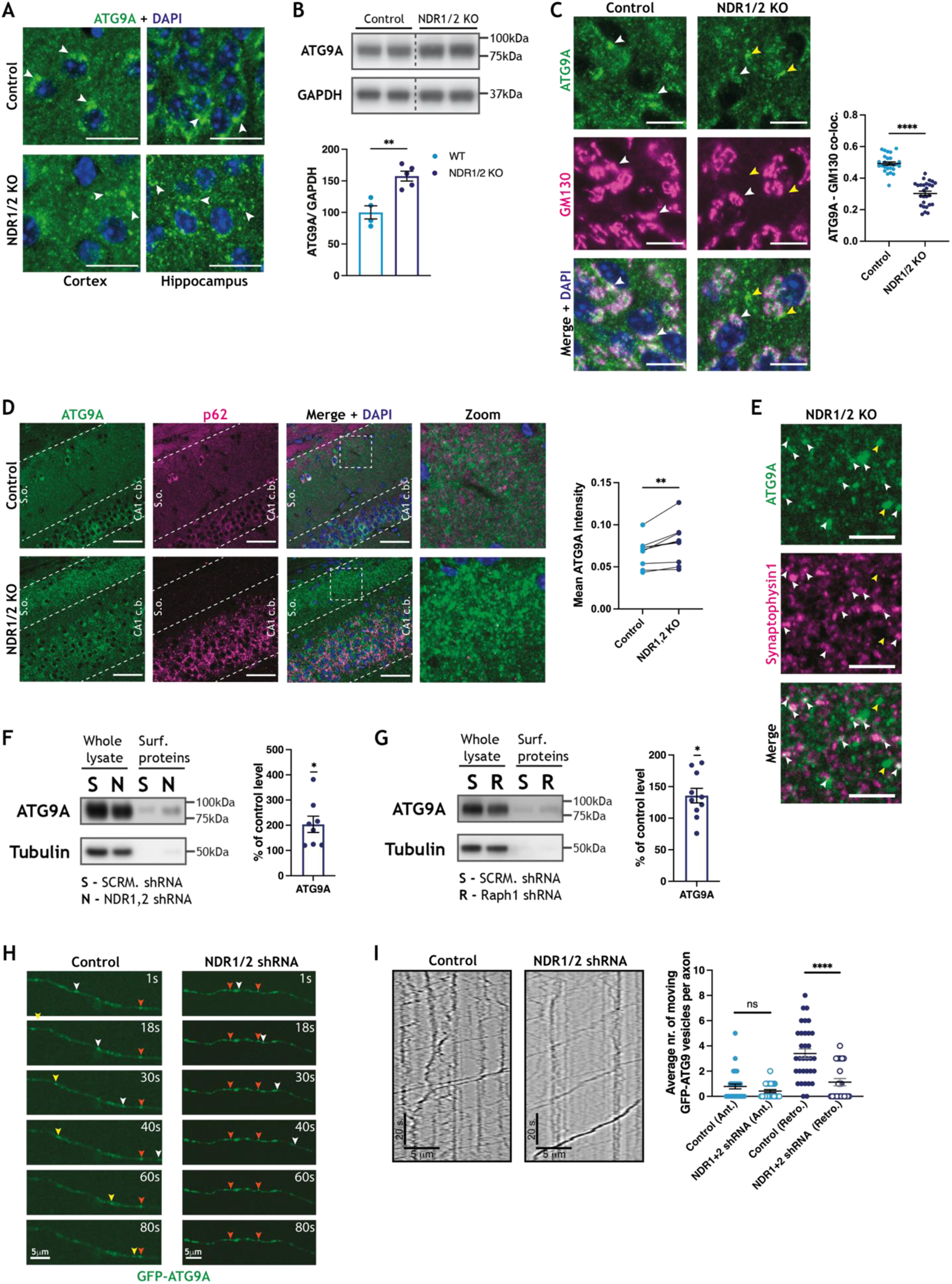
ATG9A is mislocalised in NDR1/2 KO brains. **(A)** Immunofluorescence staining of ATG9A in brain slices from 12-week-old NDR1/2 knock-out and control mice. White arrows indicate areas with increased endogenous ATG9A. Scale bars 10μm. **(B)** Western blot analyses of ATG9A in 6-week-old cortical lysates from NDR1/2 KO and control mice. GAPDH was used as a loading control. The bar graph shows quantification of ATG9A normalised against GAPDH and the data was analysed with an unpaired Student’s *t* test. n = 4-5 mice/ group. **(C)** Immunofluorescence staining of ATG9A and the Golgi marker GM130 in the CA1 area of 12-week-old mice. White arrows indicate areas where ATG9A co-localises with GM130. Yellow arrows indicate areas where ATG9A does not co-localise with GM130 in NDR1/2 knock-out mice. Scale bars 10μm. The scatter plot shows quantification of ATG9A and GM130 co-localisation expressed as Pearson correlation coefficient and the data and was analysed with a Mann Whitney test. n=28 measurements from 3 mice/ genotype. **(D)** Immunofluorescence staining of ATG9A and p62 at 12 weeks. White dotted lines delineate the stratum oriens (s.o.) area and the CA1 cell body area (CA1 c.b.). Scale bars 50μm. **(E)** Immunofluorescence stainings of ATG9A and the pre-synaptic marker synaptophysin1 in stratum oriens at 12 weeks. White arrows show ATG9A co-localising with synaptophysin and yellow arrows show ATG9A puncta that do not co-localise with synaptophysin. Scale bars 5μm. **(F-G)** Western blot analyses of surface biotinylation experiments on DIV12 rat cortical neurons infected with scramble SCRM or NDR1,2 shRNA lentivirus (B) or Raph1 shRNA (F). Surface levels of ATG9A were normalised against input. Bar graphs show surface ATG9A levels expressed as a percentage of the corresponding SCRM shRNA control level. The data was analysed using one sample *t* and Wilcoxon tests. n=6 samples/ condition from 3 independent experiments. **(H-I)** Representative images (H) and kymographs (I) of ATG9A vesicle movement in the axons of DIV11 cultured rat hippocampal neurons previously infected with a scramble shRNA lentiviral vector as a control or NDR1 and NDR2 shRNA expressing lentiviruses. Total number of moving particles were quantified from kymographs. The data was analysed using Mann-Whitney tests. n=138 mobile particles from 33 cells for scramble shRNA and n=37 mobile particles from 24 cells for NDR1/2 shRNA.

Strikingly, ATG9A clusters were observed in stratum oriens in NDR1/2 KOs, corresponding to the dendritic arbors of CA1 neurons synapsing with incoming CA3 axons (Fig. 5D, Movies 1 & 2). To test if ATG9A puncta in stratum oriens are at synapses, we co-stained ATG9A with the presynaptic marker synaptophysin 1 and found that there was a substantial colocalisation between ATG9A and synaptophysin 1 (Fig. 5E), indicating that ATG9A accumulations were partially localised near synaptic regions. Since TfR and Chl1 trafficking were altered in NDR1/2 shRNA or Raph1 shRNA infected neurons, we decided to test if ATG9A surface levels can be detected in neurons, and if these are altered when NDR1/2 or Raph1 are depleted. Indeed, surface ATG9A could be detected by surface biotinylation in primary neurons, although this represented only a very small fraction of the total ATG9A (Fig. 5F&5G). In both NDR1/2 shRNA and Raph1 shRNA infected neurons significantly more ATG9A was present at the surface compared to control neurons, confirming that ATG9A trafficking is altered (Fig. 5F&5G).

In *C. Elegans*, ATG9A endocytosis and recycling at presynaptic boutons is essential for autophagy and neuronal development (Stavoe *et al*, 2016; Yang *et al*, 2020). It is also known that autophagosomes formed at axons and presynaptic boutons travel to neuronal cell bodies for lysosomal fusion and degradation (Maday & Holzbaur, 2016; Maday *et al*, 2012). For this reason, we decided to inspect axonal ATG9A transport using live imaging in primary neurons. We used cultured hippocampus neurons infected with scramble or NDR1 and NDR2 shRNA-expressing viral vectors and transfected with GFP-ATG9A (Fig. 5H, 5I, Movies 3 & 4). In a similar fashion to autophagosomes (Maday *et al*., 2014), most ATG9A vesicles are trafficked retrogradely in axons of both control and NDR1/2-depleted cells (Fig. 5H, 5I, Movies 3&4). However, significantly less ATG9A vesicles were mobile in NDR1/2 shRNA neurons compared to the scramble shRNA condition, although net displacement and velocity were not changed (Fig. 5I & S5B, Movies 3 & 4). Furthermore, numerous stationary ATG9A clusters were observed in the axons of NDR1/2 shRNA neurons (Fig. 5H, orange arrowheads). A reduction in the number of ATG9A-positive vesicles being trafficked in axons could be a result of reduced endocytosis of ATG9A from the plasma membrane and matches the phenotype of NDR1/2 KO mice, where ATG9A accumulates in the stratum oriens. Efficient cycling of ATG9A between different membrane compartments is essential for efficient autophagy (Imai *et al*., 2016; Ivankovic *et al*, 2020; Longatti *et al*., 2012; Longatti & Tooze, 2012; Popovic & Dikic, 2014; Puri *et al*., 2013; Young *et al*., 2006), so we also assessed the trafficking transfected RFP-LC3 in the axons of NDR1/2 shRNA neurons. RFP-labelled autophagosomes predominantly travelled in the retrograde direction as previously reported (Maday *et al*., 2012). We found a similar reduction in the number of mobile retrogradely-moving autophagosomes labelled with RFP-LC3, but no changes in velocity or displacement, indicating that formation or initiation of movement, rather than transport of autophagosomes, was altered (Fig. S5C). Therefore, we postulate that the impairment in ATG9A trafficking is a major cause of defective autophagy and consequent neurodegeneration in NDR1/2 KO mice.

### Deleting Ndr1 and Ndr2 in adult mice is sufficient to replicate the phenotype of the conditional knockout mice

Since NEX expression starts at E11.5, before neuronal development is completed, we wanted to assess if the changes seen in the brains of NDR1/2 conditional knockout mice were due to a developmental defect. For this, we employed an inducible knockout mice approach using NEX-Cre^ERT2^ expressing mice (Agarwal *et al*, 2012) crossed with *Ndr1*^KO^ and *Ndr2*^flox^ mice, which were injected with tamoxifen twice a day for 5 days to induce Cre expression. To enable identification of recombined neurons the mice also express a reporter gene, Ai14-TdTomato, which switches on as soon as Cre becomes active (Madisen *et al*, 2010). Tamoxifen administration started after the mice reached adulthood, at 12 weeks of age. Following tamoxifen treatment, we aged the mice further, until 10 - 12 months of age, before analysing the brains.

As previously reported (Agarwal *et al*., 2012), Cre recombination in the cortex was partial, in a scattered ‘salt and pepper’ fashion, as observed from Ai14-TdTomato expression. In particular, recombination was low in the upper cortical layers and cortical thickness and the expression of GFAP in the cortex were not different between controls and NDR1/2 inducible KOs (iKO) (Fig. 6A& 6B). By contrast, in the hippocampus CA1-CA3 areas Cre recombination efficiency was close to 100% (Fig. 6A), as expected (Agarwal *et al*., 2012). In NDR1/2 iKOs, the cell body layering of CA1 was disrupted, stratum radiatum was smaller and GFAP staining was highly increased in the hippocampus (Fig. 6A& 6B), similarly to the NEX-Cre conditional NDR1/2 KO mice. These results indicate that deletion of NDR2 in NDR1 knockout adult mice leads to loss of neurophil and degeneration in hippocampus, while it is possible that the lack of changes in the cortex of NDR1/2 iKO mice is due to the low recombination rate.

**Figure 6.**
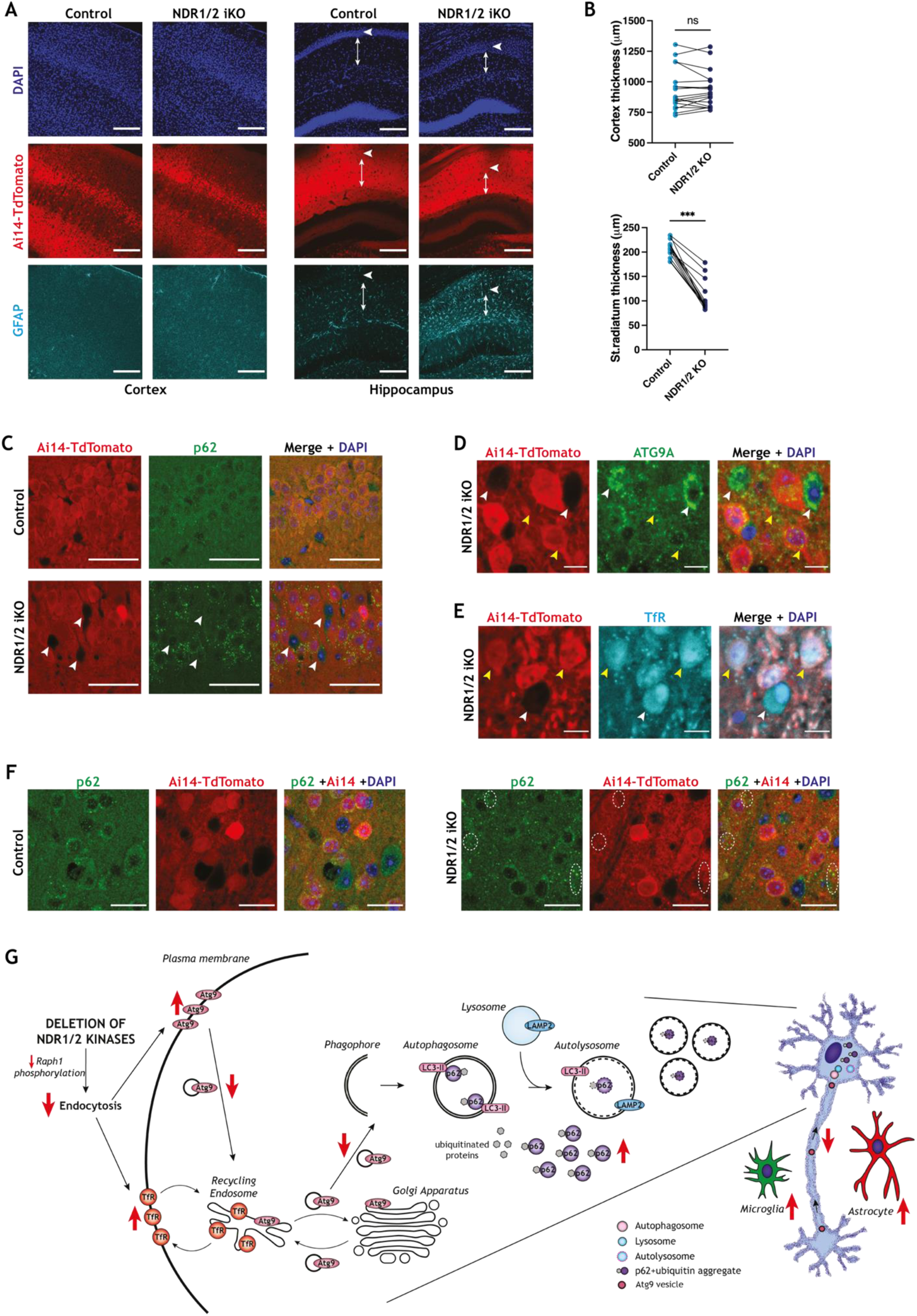
Dual loss of NDR1/2 in adults is sufficient to induce p62 accumulation, ATG9A mislocalisation and neuropil loss. **(A)** Immunofluorescence staining of GFAP in the brains of NDR1/2 inducible knock-out (NDR1/2 iKO) and control mice. White arrows show the distribution of neurons within the CA1 cell body layer. Arrowed lines show the length of stratum rediatum. Scale bars 200μm. **(B)** Graphs showing quantifications of cortex and stratum radiatum thickness. The data was analysed using paired Student’s *t* tests, n = 18 measurements from 3 mice/ genotype. **(C)** Immunofluorescence staining of p62 in NDR1/2 iKO and control mice in the CA1 area. White arrows show Ai14-TdTomato negative cells, which have not been recombined and lack p62 accumulations and scale bars represent 50μm. **(D-E)** Immunofluorescence staining of ATG9A (D) and transferrin receptor (TfR) (E) in the brains of NDR1/2 iKO and control mice in the CA1 area. White arrows show Ai14-TdTomato-negative cells, which have a perinuclear distribution of ATG9A (D) and lack TfR accumulations (E). Yellow arrows show Ai14-TdTomato-positive cells, which exhibit a punctate distribution of ATG9A (D) and a high number of TfR puncta (E). Scale bars 10μm. **(F)** Immunofluorescence staining of p62 in areas of the cortex with high Ai14-TdTomato expression. White circles highlight areas with p62 accumulations in NDR1/2 iKO brains. Scale bars are 20μm. **(G)** Schematic diagram depicting membrane trafficking events affected by NDR1/2 in neurons. Loss of NDR1/2 kinases downregulates endocytosis, likely via reduction in phosphorylation of Raph1/ Lpd. As a result, surface levels of ATG9A and TfR are increased. Altered endocytosis of ATG9A causes reduced axonal retrograde ATG9A transport and reduced cell body/Golgi localisation of ATG9A. ATG9A mislocalisation interferes with autophagosome formation, leading to progressive accumulation of p62-ubiquitinated proteins.

Despite a very efficient recombination in the hippocampus, not all the neurons in the CA1 area had reporter expression (Fig. 6C). This enabled us to compare side by side neurons that do not express NDR kinases and neurons expressing only NDR2. p62 immunofluorescence stainings revealed p62 accumulations in the CA1 area of NDR1/2 iKO mice in Ai14-TdTomato positive cells (Fig. 6C - yellow arrows), while Ai14-TdTomato negative neurons lacked such p62 accumulations (Fig. 6C - white arrows). Similar results were observed with ATG9A and TfR in CA1 neurons. Ai14-TdTomato negative cells had a perinuclear ATG9A distribution, as seen in control animals, and lacked TfR puncta (Fig. 6D &6E – white arrows). By contrast, in Ai14-TdTomato positive neurons less ATG9A was found in the perinuclear area, but additional ATG9A puncta had a dispersed distribution and TfR accumulations seemed concentrated in these cells (Fig. 6D &6E – yellow arrows). Finally, immunofluorescence assessments from areas of the cortex with the highest recombination rate revealed that p62 puncta accumulated in NDR1/2 iKO mice (Fig. 6F – white dotted circles), but this was at a lesser extent than the hippocampus, in accordance with lower recombination levels. In conclusion, these results showed that deleting NDR kinases in adult neurons is sufficient to replicate the TfR trafficking deficits, ATG9A mislocalisation and p62 accumulation phenotypes observed in NDR1/2 conditional knockout brains. Importantly, these results indicate that the roles of NDR1/2 in neuronal homeostasis are not due to their function during neuronal development.

Our results are consistent with a model in which NDR1/2 kinases regulate neuronal endocytosis and membrane trafficking via multiple substrates, one of which is the endocytic protein Raph1. Upon loss of NDR1/2 TfR cycling is changed and ATG9A surface levels, neuronal localisation and its axonal transport are highly altered, highlighting a defect in ATG9A trafficking. We propose that the impairment in ATG9A localisation and trafficking contributes to reduced autophagy and the subsequent accumulation of p62 and ubiquitinated proteins that is observed in NDR1/2 KO mice (Fig. 6G). It is worth mentioning that other NDR1/2 substrates and downstream signalling events could also impair autophagy and protein homeostasis in parallel.

## Discussion

In this study we report a novel function of NDR1 and NDR2 in neuronal protein homeostasis and autophagy, which is critical for preserving neuronal health. NDR1 and NDR2 have been implicated in YAP1 regulation and tumourigenesis (Cornils *et al*., 2010; Zhang *et al*, 2015). NEX-Cre mediated deletion of the other two members of NDR kinase family, LATS1 and LATS2, causes YAP1 activation and tumour formation in brain (Eder *et al*, 2020). However, we did not find any brain tumours in NDR1/2 KOs. Thus, our study shows that NDR1 and NDR2 do not regulate YAP1 in NEX-positive neural precursors. While YAP1 phosphorylation was unchanged, NDR1/2 KO brains had significant increases in proteins associated with human neurodegenerative diseases (e.g. ApoE, Htt, APP, PRNP), highlighting similarities between this mouse model and human disorders. Furthermore, *Ndr*2 loss-of-function mutations are found in canine early retinal degeneration (Goldstein *et al*, 2010) and loss of *Ndr1* leads to compromised homeostasis in mouse retina (Leger *et al*., 2018). Together with our findings this indicates that NDR kinases could be novel targets for protection against neurodegeneration.

To gain mechanistic insight into NDR1/2 signalling *in vivo*, we used quantitative proteomics approaches in double knockout mice. We reveal NDR1/2 kinases as novel regulators of neuronal protein homeostasis. Our TMT based quantitative proteomics experiments were done at P20, when there was no difference in hippocampus GFAP levels between controls and double KOs, to allow us to identify the earliest changes directly downstream of NDR kinases. The phosphoproteome analysis did not reveal a direct autophagy protein as an NDR1/2 substrate candidate, although PI4KB has been previously implicated in autophagy (Judith *et al*., 2019). Instead NDR1/2 phosphorylates a repertoire of membrane trafficking regulators. Using *in vitro* kinase assays, we validated one of the top candidates we identified, Raph1, as a novel NDR1/2 substrate. The data presented here is of critical value for understanding the neuronal functions and downstream effectors NDR1/2 kinases.

Given the similarities between our NDR1/2 KO model and brain-specific autophagy defective mouse models (Hara *et al*., 2006; Komatsu *et al*., 2006), we hypothesised that autophagy may be affected. We show that together NDR1 and NDR2 are essential regulators of autophagosome formation in neurons *in vivo*. Reduced numbers of autophagosomes in neuronal cell bodies and increased overall p62 in NDR1/2 KO brains, as well as reduced LC3 trafficking in axons and reduced LC3 lipidation in primary neurons expressing NDR1/2 shRNA, collectively indicate that autophagosome formation and degradation could both be affected in neurons that lack NDR1 and NDR2. Our results agree with NDR1’s reported positive roles in autophagy (Joffre *et al*., 2015; Martin *et al*., 2019). Interestingly, while this manuscript was being prepared a recent publication identified the *Drosophila* NDR kinase trc as a binding partner of LC3 (Tsapras *et al*, 2022). Previously, NDR1 was shown to enhance autophagy by regulating the interaction of Beclin1 with its effectors (Joffre *et al*., 2015) and by increasing Exportin-1 (Xpo1) activity, leading to increased Beclin1 levels in the cytoplasm (Martin *et al*., 2019). In our NDR1/2 KO mouse model, total protein levels and the phosphorylations detected on Xpo1 or Beclin1 were not altered (Supplemental Data 1 & 5). Although we tried to account for the presence of activated glia in KO brains, some neuron-specific proteomic differences could still go undetected.

The autophagy pathway relies heavily on efficient membrane trafficking (Judith *et al*., 2019; Lamb *et al*., 2013; Puri *et al*, 2020; Puri *et al*., 2013). Considering the roles of NDR1/2 substrates identified in this paper and previously (Ultanir *et al*., 2012), we hypothesised that NDR kinases may play a role in autophagy via their involvement in membrane trafficking and endocytosis. We showed pronounced effects on TfR trafficking, in particular transferrin endocytosis, when NDR1/2 or their novel substrate Raph1 are knocked down in neurons. Raph1/Lpd interacts with and functions via the actin regulators, Ena/VASP and the Scar/Wave-Arp2/3 complexes, to mediate effective cell migration and neuronal morphogenesis (Krause *et al*., 2004; Law *et al*, 2013; Michael *et al*, 2010; Sundararajan *et al*, 2019). Additionally, Raph1/Lpd is required for CME and fast endophilin-mediated endocytosis (Boucrot *et al*., 2015; Vehlow *et al*., 2013) (Chan Wah Hak *et al*., 2018), explaining the effect of Raph1 shRNA on transferrin uptake. However, it is worth mentioning that this seems to be a neuron-specific function, since in HeLa cells Lpd knockdown impaired receptor induced CME uptake of EGFR but not constitutive TfR CME (Vehlow *et al*., 2013). Interestingly, like NDR kinases Lpd is also required for neuronal dendritic arborisation (Tasaka *et al*., 2012), suggesting that a common role in endocytosis and autophagy, as observed here, may contribute to this function.

Considering the strong impact of NDR1/2 loss on membrane trafficking, we next hypothesised that NDR1/2 loss could affect the trafficking and localisation of the only transmembrane autophagy protein, ATG9A. ShRNA-mediated depletion of NDR1 and NDR2 resulted in increased ATG9A surface levels, indicating that ATG9A endocytosis could also be affected by NDR1/2, perhaps owing to co-trafficking of ATG9A and TfR (Longatti *et al*., 2012; Longatti & Tooze, 2012; Puri *et al*., 2013). ATG9A is known to be endocytosed from the plasma membrane via CME and when endocytosis is inhibited with dynasore, dominant-negative dynamin or siRNA targeting AP2, ATG9A disperses from Golgi and accumulates in a different compartment close to the plasma membrane (Puri *et al*., 2013; Puri *et al*, 2014). In NDR1/2 KO brains, both TfR and ATG9A formed punctate accumulations. In *C. elegans* neurons, ATG9A is present at synaptic sites and is essential for autophagy at synapses (Stavoe *et al*., 2016; Yang *et al*., 2020). Inhibition of endocytosis in these neurons also caused an accumulation of ATG9A in discrete locations near the synaptic sites, where it colocalised with clathrin (Yang *et al*., 2020). In NDR1/2 KO mouse brains, ATG9A severely accumulated in peripheral regions of neurons, where it often colocalised with the synaptic marker, synaptophysin 1. In addition, neurons expressing NDR1/2 shRNA exhibited fewer GFP-ATG9A positive vesicles trafficking retrogradely in axons, in agreement with reduced endocytosis of ATG9A in axon boutons. ATG9A’s cycling between endosomal compartments and Golgi is known to be critical for starvation-induced autophagy in dividing cells, as well as constitutive autophagy in neurons (Imai *et al*., 2016; Ivankovic *et al*., 2020; Judith *et al*., 2019; Orsi *et al*., 2012; Popovic & Dikic, 2014; Puri *et al*., 2020; Puri *et al*., 2013; Young *et al*., 2006). Given the clear ATG9A mislocalisation and neurodegeneration phenotypes exhibited by NDR1/2 knockouts, we attribute the autophagy deficit in these mice to impaired ATG9A function. Interestingly, altered ATG9A localisation can be caused by the Parkinson disease (PD)-associated VPS35 mutation D620N, which also impairs autophagy (Zavodszky *et al*, 2014a; Zavodszky *et al*, 2014b). Furthermore, Parkin loss-of-function mutations also cause ATG9A mislocalisation downstream of VPS35 (Williams *et al*, 2018), collectively highlighting that efficient ATG9A trafficking is important to prevent neurodegeneration. Our findings significantly expand our understanding of the cellular and *in vivo* roles of NDR1/2 kinases in mammalian neurons.

## Materials and methods

### Mouse handling and mouse lines

All mouse handling was performed according to the regulations of the Animal (Scientific Procedures) Act 1986. Mice were housed in an animal facility of the Francis Crick Institute on an alternating 12-hour light-dark cycle with free access to food and water. All lines are on a C57BL/6J background. Both male and female mice were used and randomly allocated to experimental groups according to genotypes. As much as possible comparisons were made between littermates with different genotypes.

NEX-Cre (*Neurod6^tm1(cre)Kan^*, MGI:2668659) and Nex-CreERT2 (*Neurod6^tm2.1(cre/ERT2)Kan^*) mice were a gift from Dr. Klaus Nave and Dr. Markus Schwab. *Ndr1*^KO/KO^ (B6;129P2-*Stk38^tm1/FMI^*) and *Ndr2*^flox/flox^ (B6CF2;129P2-*Stk38l^tm3/BAH-FMI^*) mice were provided by Dr. Brian Hemming. Ai14 (B6;129S6-*Gt(ROSA)26Sor^tm14(CAG-tdtomato)Hze^*/J; Stock No: 007908) and Thy1-YFP lines (B6.Cg-Tg(Thy1-YFP)HJrs/J; Stock No: 003782) were purchased from Jackson Laboratories.

For the *Ndr2* inducible knockout approach, *Ndr1*^KO^*Ndr2^f^*^lox^ mice were crossed with mice expressing *Nex*^CreERT2^. After the mice reached adulthood, at 12 weeks of age, they received a 5-day course of Tamoxifen (Sigma) in corn oil - 1mg, twice daily - to induce Cre expression. Following the Tamoxifen injections, the mice were weighed weekly and aged further until at least 30 weeks of age, before harvesting of the brain.

For western blot analyses brains were harvested at P20 or 6 weeks of age. The mice were culled by cervical dislocation, the brain was removed from the skull and the hippocampus or cortex were dissected and flash frozen in liquid nitrogen. For immunofluorescence P20, 12-week and 20-week-old animals were transcardially perfused with 4% ice-cold paraformaldehyde (PFA) under terminal anaesthesia prior to brain harvesting and post-fixation in 4% PFA overnight.

### Immunofluorescence, histology and imaging

50μm thick coronal slices were obtained from the fixed brains with a Leica VT1000 S vibrating blade microtome (Leica) and used for immunofluorescence staining. The slices were blocked in buffer containing 10% serum and 0.02% Triton-X in PBS and then incubated in primary antibodies over-night at 4°C. For the following antibodies a 20min antigen retrieval step in citrate buffer (10 mM Sodium citrate, 0.05% Tween 20, pH 6.0) at 95°C was carried out prior to blocking: ubiquitin, transferrin receptor, ATG9A, GM130 and VPS35. Secondary antibody incubation was 1h at room temperature and nuclei were stained using DAPI. Sections were mounted on slides with Fluoromount. All images were acquired with a Zeiss Invert880 confocal microscope. Images obtained using the 4X (Fig. 1E) and 10X (Fig. 1C, 1D, 1H, S1A, 2A, 4A, 6A) objectives were acquired as single images, while the rest of the images were acquired with the 40X or 63X objective as Z stacks (1μm interval) and 3 slices were maximally projected to obtain the images used in the figure panels.

For histological analysis fixed brains were dehydrated and embedded in paraffin after completion of 4% PFA fixation. 4μm sections were prepared and stained with H&E. Images were acquired using Olympus VS120 Slide Scanner.

### Primary neuronal cell culture

Rat cortical and hippocampal neurons were cultured from E16.5 embryos from a wild-type Long Evans mother. All the hippocampi and all the cortices from one litter were dissected and all tissue from the sain brain area was pooled together. Dissociation of neurons was carried out by incubation with 0.25% trypsin (Gibco) for 20 minutes at 37°C, followed by Hank’s Balanced Salt Solution (Gibco) washes and resuspension. The cells were then counted and plated on previously coated surfaces in plating media. The following densities were used for plating: 200,000 cells per 18mm glass coverslip, 300,000 cells per 22mm well of a 12-well plate and 500,000 cells per 35mm glass-bottom dishes or per 35mm plastic dishes. The plating media contained 10% fetal bovine serum (FBS), 0.5% dextrose, sodium pyruvate (Gibco), 2 mM glutamine (Gibco) and penicillin/streptomycin in minimum essential medium (Gibco). The coating on the glass coverslips, dishes and plates had been applied overnight at 37°C and contained 60μg/ml poly-D-lysine (Sigma) and 2.5μg/ml laminin (Sigma) in 0.1M borate buffer containing. After approximately 4 hours the neurons were transferred to maintenance media containing B27 (Sigma), 0.5mM glutamine, 12.5μM glutamate, penicillin/streptomycin and ciprofloxacin in neurobasal medium (Gibco). The neurons were kept at 37°C and 5% CO_2_ until use and media change was carried out every 3-4 days, by replacing about 1/3 of the media with fresh maintenance media.

### HEK293T cell culture

HEK293T cells were kept in Dulbecco’s Modified Eagle Medium (Gibco) supplemented with 10% FBS and 1% penicillin/streptomycin. Transfections were carried out using Xtremegene 9 (Roche) according to manufacturer’s protocol. For the cells treated with 100nM Bafilomycin A1 (Sigma) for 4h the treatment and subsequent lysis for western blot analysis happened 48h after transfection. The HEK293T cells used to purify Raph1 for the kinase assay were lysed 72h after transfection.

### Western blotting

Flash frozen mouse brain areas were homogenised by sonication in 2X sample buffer (Pierce™ LDS Sample Buffer, Thermofisher) containing 0.2M DTT. Lysates were then centrifuged at 13,000 g for 15 minutes, supernatants removed and denatured at 70°C for 10 minutes. Neurons and HEK293T cells treated with Bafilomycin A1 (Sigma) were lysed directly in 2X sample buffer containing 0.2M DTT, followed by sonication and incubation at 70°C for 10 minutes. All lysed samples were run on NuPAGE 4–12% Bis-Tris polyacrylamide gels (ThermoFisher) and transferred to a polyvinylidene difluoride membrane (Millipore) using wet transfer. After blocking in 5% non-fat milk in TBST for 1h, the membranes were incubated with primary antibodies overnight at 4°C or at RT for 1-2h. Peroxidase-conjugated (HRP) secondary antibody incubation was carried out at RT for 1h. For western blot detection the membranes were incubated with ECL (Amersham ECL Prime Western Blotting Detection Reagent) and visualised with a chemiluminescence digital imaging system (Amersham Imager 600RGB) or developed using film (Amersham Hyperfilm ECL; western blots in Fig. S1B only).

### Protein purification

Constitutively active analogue sensitive NDR1 (NDR1 CA MA) and constitutively active kinase dead NDR1 (NDR1 CA KD) were purified as detailed in Ultanir et al., 2012.

Raph1-WT, Raph1-S192A, Raph1-S571A and Raph1-SAx2 (S192A & S571A) were purified from HEK293T cells using the pCAG-DEST-3xFLAG-2xStrep expression vector. 72h after transfection the cells were washed once with cold PBS and lysed in buffer containing 50 mM Tris–HCl pH 8.0, 5% glycerol, 150 mM NaCl, 10mM MgCl_2_, 0.1% Triton X-100, 1x protease inhibitor cocktail (Roche) and 1:5000 Pierce Universal Nuclease (ThermoFisher). The lysates were incubated at 4°C for 30min, rotating, to allow for solubilisation and then centrifuged at 21,000 g for 15 min at 4°C. To immunoprecipitate FLAG-tagged proteins the supernatant was incubated with anti-FLAG M2 Affinity Gel (Sigma) for 1h at 4°C, rotating. Proteins bound on FLAG beads were washed once with lysis buffer and once with a wash buffer containing 50 mM Tris–HCl pH 8.0, 150 mM NaCl, 10mM MgCl_2_ and 1x protease inhibitor cocktail. Raph1/Lpd is heavily phosphorylated under normal conditions, so the bead-bound proteins were dephosphorylated in wash buffer containing 1:50 Lambda Protein Phosphatase (New England Biolabs) and 1 mM MnCl_2_ to make the phosphosites more accessible for a kinase assay. The dephosphorylation was carried out at room temperature for 30min and the samples were kept rotating. The bead-bound proteins were then loaded onto pre-washed Pierce™ Centrifuge Columns (ThermoFisher) and washed twice with high salt wash buffer (500mM NaCl) and twice with normal salt wash buffer. Bound proteins were eluted in wash buffer containing 100µg/ ml 3xFLAG peptide.

### *In vitro* kinase assays

To establish the concentration of the proteins needed for kinase assays purified proteins were separated via gel electrophoresis together with known amounts of BSA. The gel was stained with Coomassie stain and imaged using colorimetric assessment on a digital imaging system (Amersham Imager 600RGB). The intensity of the bands was quantified and the concertation of Raph1 and NDR1 constructs established based on the known amounts of BSA loaded on the same gel.

For the kinase assay 500ng Raph1 was incubated with 50ng NDR1 in buffer containing 20mM Tris–HCl pH 7.5, 10mM MgCl_2_, lµM okadaic acid, 1mM DTT, 1x protease inhibitor cocktail (Roche),100µM ATP and 0.5 mM 6-benzyl-ATPcS (Biolog) at 30°C, rotating, for 30min. The reactions were quenched with 20mM EDTA and alkylated with 5mM p-nitrobenzyl mesylate (Abcam) for 30min at room temperature. The proteins were solubilised in sample buffer (Pierce™ LDS Sample Buffer, Thermofisher) containing 0.1M DTT and denatured by incubation at 70°C for 10min, in preparation for western blot analysis.

### Transferrin recycling assay

Rat hippocampal neurons were grown on coverslips and transfected with an empty EGFP-expressing plasmid (pLL3.7) at DIV6. 4h later the cells were transduced with lentiviruses expressing shRNAs. At DIV11, the neurons were starved in DMEM (Gibco) with added penicillin/streptomycin for 30 min at 37°C and 5% CO_2_ and then incubated with 50 μg/ml transferrin-Alexa-568 (ThermoFisher) in complete culture media for 20 min at 4°C. Next, the neurons were returned to 37°C and 5% CO_2_ for 20 min to allow transferrin uptake (pulse). At the end of the pulse, the cells were washed three times with PBS and reincubated in culture media devoid of labelled transferrin for 0, 20, or 60 min (chase). Following chase neurons were washed once more with PBS and fixed with 4% PFA and 4% sucrose in PBS. After mounting of coverlips with Fluromount, images were acquired with a Zeiss Invert880 confocal microscope.

### Biotinylation of surface proteins

Rat cortex neurons were grown on 35mm dishes and transduced with lentiviruses expressing shRNAs at DIV6. The protocol for the biotinylation assay was adapted from (Twelvetrees *et al*, 2010). At DIV12, the neurons were washed with cold PBS (all wash steps were carried out with PBS supplemented with 1 mM CaCl_2_ and 0.5 mM MgCl_2_) and incubated with 1 mg/ml biotin (EZ-Link™ Sulfo-NHS-Biotin, ThermoFisher), shaking, at 4°C for 12 min. The following steps, prior to cell lysis, were all carried out on ice, using cold solutions. After biotin incubation the neurons were washed with PBS twice and the biotinylation reaction was quenched with a buffer containing 1mg/ml BSA in PBS. After three more washes with PBS the cells were lysed in RIPA buffer (ThermoFisher) and pelleted by centrifugation at 13,000g for 15 minutes in a refrigerated centrifuge (at 4°C). Part of each supernatant was saved as input (total lysate fraction) and mixed with equal volumes of 2X sample buffer, prior to incubation at 70°C for 10 minutes, in preparation for western blot analysis. The rest of the supernatants were incubated with NutrAvidin beads (ThermoFisher) for 2h at 4°C on a rotator. After three washes with RIPA buffer the proteins were eluted off the beads in 2X sample buffer and incubated at 70°C for 10 minutes, in preparation for western blot analysis.

### Live imaging

Rat hippocampal neurons were grown on 35 mm glass-bottom dishes and transfected with GFP-ATG9A or RFP-LC3 expressing plasmids at DIV6. 4h later the cells were transduced with lentiviruses expressing shRNAs. On DIV11, glass bottoms were placed in the pre-warmed chamber (37°C, 5% CO_2_) of a NIKON CSU-W1 Ti2 Confocal spinning disk microscope with s Dual Prime 95B camera. A 100x (oil) objective was used to identify axons and axonal imaging was carried out within 200 μM of the soma. Two-minute videos were acquired at 2 frames/ second. Approximately 25 μM of each imaged axonal segment (15-line width) was selected to generate kymographs using the KymographClear plugin (Mangeol *et al*, 2016) in FIJI. The kymographs were then filtered (DoG filter with a pixel radius ratio of 2:0.5 - signal to noise) and resized so that they all have the same size. Tracing and motility analysis were carried out with the KymoAnalyzer plugin (Neumann *et al*, 2017) in FIJI. In terms of tracing, only the objects that were present in the majority of frames were manually traced. For the generation of displacement and velocity (average) data, the following conversion values were used: pixel size - 0.11 μm, frames per second - 2.

### Antibodies, plasmids and viral vectors

The following primary antibodies were used for immunofluorescence: mouse GFAP (1:2000, Sigma G6171), rat Ctip2 (1:500, Abcam ab18465), rabbit Iba1 (1: 200, Wako Chemicals 019-19741), guinea-pig p62 (1:400, Progen GP62-C), rabbit LC3-B (1:200, Cell Signalling 3868), rabbit ubiquitin (1:100, Abcam ab134953), rat LAMP2 (1:300, Abcam ab134953), mouse transferrin receptor – TfR (1:250, ThermoFisher 13-6800), rabbit ATG9A (1:200, Abcam ab108338), mouse GM130 (1:200, BD 610822), rabbit VPS35 (1:200, Abcam ab157220), mouse Synaptophysin1 (1:500, Synaptic Systems 101 011), goat TdTomato (1:400, SICGEN AB8181-200). All fluorescent secondary antibodies were purchased from Jackson ImmunoResearch and used at 1:500 dilution.

The above mentioned GFAP, p62, ubiquitin, TfR, ATG9A were also used for western blotting at 1:1000 dilution, in addition to the following antibodies: mouse GAPDH (1:10000, Abcam ab8245), mouse α-tubulin (1:20000, Sigma T9026), mouse NDR1 (1:1000, Sigma SAB1408832), rabbit NDR2 (1:500, previously published in Ultanir et al., 2012), mouse PSD95 (1:1000, ThermoFisher MA1-045), mouse Homer1 (1:1000, Synaptic Systems 160011), mouse GluR1 (1:1000, Millipore MAB2263), rabbit LC3-B (1:2000, Abcam ab48394), rabbit Chl1 (1:1000, Abcam ab106269), rabbit Raph1 (1:1000, Abcam ab121619), rabbit thiophosphate ester (1:200, Abcam ab92570). The Raph1 phospho-S192 antibody is a custom-made rabbit polyclonal antibody raised against the following Raph1 peptide: C-TNQHRRTAS*AGTVScoNH_2_ (Covalab). All HRP-conjugated secondary antibodies were purchased from Jackson ImmunoResearch and used at 1:10000 dilution.

The following plasmids were used for transfection: NDR1-CA-HA, NDR1-KD-HA and NDR1-WT-HA (previously published in Ultanir et al., 2012), GFP-ATG9A (gift from Sharon Tooze), RFP-LC3 (gift from Max Gutierrez), Raph1-WT-GFP and Raph1-S192A-GFP (generated via site-directed mutagenesis using the Raph1-WT-GFP plasmid). The pCAG-DEST-Raph1-WT-3xFLAG-2xStrep was generated via Gateway cloning according to the manufacturer’s protocol (ThermoFisher) starting from a pENTR3C-Raph1-WT plasmid and a pCAG-DEST-3xFLAG-2xStrep which was generated by HIFI assembly (NEB) using pCAG-EGFP (Addgene 89684), and a geneblock (IDT) harboring a TEV site, a 3xFLAG, a 6x Glycine linker and a Twin-Strep tag. S192, S571 or both sites were then mutated to alanines using site-directed mutagenesis to obtain phospho-mutant Raph1 constructs.

The shRNA-expressing viral vectors were acquired from VectorBuilder. NDR2 shRNA and Raph1 shRNA were ordered from the VectorBuilder database (rStk38l[shRNA#2] for NDR2 and rRaph1[shRNA#2] for Raph1) and the sequence for NDR1 shRNA has been previously published in Ultanir et al., 2012.

### Image analysis

Image analysis was carried out using CellProfiler.

The thickness of the cortex and the length of stratum radiatum were measured in DAPI stained slices from 3 different mice/ genotype. The slices were matched between controls and double knockout animals in terms of the relative area of the brain that they belonged to. A straight line was drawn to measure the desired length and the same angle was kept for this line between matching slices.

For the measurement of transferrin-568 signal in the transferrin recycling assay one image/ neuron was obtained using the 63X objective and 2X zoom, by opening the pinhole and establishing optimal focus. The EGFP cell fill was used to delineate the cell body area and transferrin signal was measured only in the cell body and normalised against the surface area.

For the measurement of p62 and LC3 puncta from brain slices Z stacks were acquired with the 63X objective, 2X zoom, 1μm interval between slices, to cover the entire volume of neurons. YFP and DAPI signals were used to delineate the cytoplasm area and the number of puncta were measured only in the cytoplasm area. For each neuron the number of puncta from each slice was normalised against the surface area of the cytoplasm and then added together to obtain the total puncta/ cell.

Co-localisation assessments between p62 and ubiquitin and ATG9A and GM130 were carried out using Z stacks acquired with the 63X objective, 3X zoom and 2μm interval between slices. The DAPI signal was used to identify nuclei and the nuclear areas where discounted when assessing the intensity of the above-mentioned markers. Co-localisation between themarkers was measured using the Pearson correlation coefficient.

ATG9A signal in the stratum oriens area was measured in Z stacks acquired with the 40x objective to have the entire width of stratum oriens area within the frame (similar to images in Fig. 5D). The Z stacks, comprising of 10 slices acquired with a 1μm interval, were projected as an average and the intensity of the ATG9A signal was measured from the average projection.

### Transmission electron microscopy (TEM)

For transmission electron microscopy (TEM), mouse brains were perfusion fixed in 4% PFA and 50µm sections were obtained by vibratome sectioning. The slices to be analysed were stored in 0.1M PB until further processing. The slices were then transferred to polypropylene 24-well plates (Caplugs Evergreen, Buffalo, USA) and processed using a Pelco BioWave Pro+ microwave (Ted Pella Inc, Redding, USA) and following a protocol adapted from the National Centre for Microscopy and Imaging Research protocol (Deerinck *et al*, 2010), which was detailed in (Eder *et al*., 2020). After processing and embedding of slices into Durcupan ACM® resin (Sigma), the blocks were trimmed to a trapezoid around the hippocampus CA1 area. The samples were then sectioned using a UC7 ultramicrotome (Leica Microsystems, Vienna, Austria) and 70nm sections were picked up on Formvar-coated G50HEX copper grids (Gilder Grids Ltd., Grantham, UK). The sections were viewed using a 120 kV Tecnai G2 Spirit transmission electron microscope (FEI Company, Eindhoven, Netherlands) and images were captured using an Orius CCD camera (Gatan Inc., Pleasanton, USA). 22 control cells and 16 NDR1/2 knockout cells from respectively 6 and 3 different areas of the CA1 region were imaged at various magnifications. 16500x images were randomised and anonymised using Advanced Renamer, for unbiased analysis of the mitochondria morphology.

### Mass spectrometry and data processing

Liquid chromatography-tandem mass spectrometry (LC-MS/MS) was used to analyse differences in the total proteome and the phospho-proteome of P20 NDR1/2 KO mice compared to control animals. The hippocampus was dissected from 5 control mice and 5 NDR1/2 KO mice and snap frozen in liquid nitrogen prior to lysis and labelling with a TMT 10-plex reagent. Sample preparation, data acquisition and mass spectrometry analysis were carried out as previously described in (Eder *et al*., 2020) with some modifications. Briefly, pooled TMT-labelled (Lot# UK288606) samples were cleaned up using a C_18_ Sep Pak Vac 1cc, 50 mg (Waters) and dried. The peptide mixture was then subjected to high-select sequential enrichment of metal oxide affinity chromatography (SMOAC) to capture phospho-peptides. It was first passed through a high-select TiO_2_ phospho-enrichment column (Thermo Scientific, A32993) following manufacturer protocol. Flow-through and wash fractions were combined, dried and subsequently used for Fe-NTA phospho-enrichment (Thermo Scientific, A32992). One tenth of the combined flow-through and wash fractions from this enrichment was used for total proteome analysis. The eluates from SMOAC were freeze-dried, solubilised and pooled together. Total proteome and phospho-proteome samples were subjected to high pH reversed phase fractionation (Thermo Scientific, 84868), dried and resolubilised in 0.1% TFA prior to LC-MS/MS. The samples were then subjected to HCD MS2 and MSA SPS MS3 fragmentation methods as described in Jiang et al. Data were acquired in data-dependent acquisition (DDA) mode using Orbitrap Eclipse Tribrid mass spectrometer (Thermo Scientific). Acquired data were processed using MaxQuant v 2.0.3.0. Processed data were then analysed using an R-coding script tailored to isobaric labelling mass spectrometry. The script was generated as a hybrid using the backbone and differential gene expression analysis of ProteoViz package (Storey *et al*, 2020) as a general script workflow and borrowing the normalisation script from Proteus package (Gierlinski *et al*, 2018). Briefly, the “proteinGroups.txt” and “Phospho (STY)Sites.txt” tables were read into matrices and filtered for “reverse” hits, “potential contaminant” and proteins “only identified by site”. Data were then normalised using CONSTANd (Maes *et al*, 2016), log2-transformed and differentially analysed using Linear Models for Microarray Data (LIMMA). Significant hits were called proteins and phosphopeptides with p-value < 0.005. Volcano plots were generated using ggrepel (a ggplot2 extension) as part of the tidyverse. Scripts provided in Supplemental data.

### Statistical analysis

Statistical analysis was performed using GraphPad Prism 7 software with the statistical tests indicated in the figure legends. Statistical significance was observed as: ns = not significant (p > 0.05), * = p < 0.05, ** = p < 0.01, *** = p < 0.001, **** = p< 0.0001. All error bars represent SEM.

Where not quantified western blots shown are representative of experiments using at least 3 mice/ genotype or repeated in 3 independent cellular culture experiments. All immunostaining experiments have been repeated in at least 3 mice/ genotype and representative images have been chosen for figures

## Acknowledgments

We thank all members of the Ultanir lab for valuable discussions and technical help. We thank Dr Maximiliano Gutierrez for his input and guidance. We thank Emma Nye for providing expertise in histopathology.

## Funding

This research is funded by The Francis Crick Institute, which receives its core funding from Cancer Research UK (FC001201), the UK Medical Research Council (FC001201), and the Wellcome Trust (FC001201). FR is funded by a BBSRC-GSK CASE PhD studentship.

## Author contributions

Conceptualization: SKU, FR

Technical assistance: SC

Data collection and analysis: FR, SRM, NE, AM, M-CD,

Supervision: SKU, SAT, JLS, HRF, APS, LC

Writing—original draft: SKU, FR

Writing—review & editing: all authors

## Conflict of interest

The authors declare that they have no conflict of interest.

## Supplemental Figure Legends

**Figure S1.**
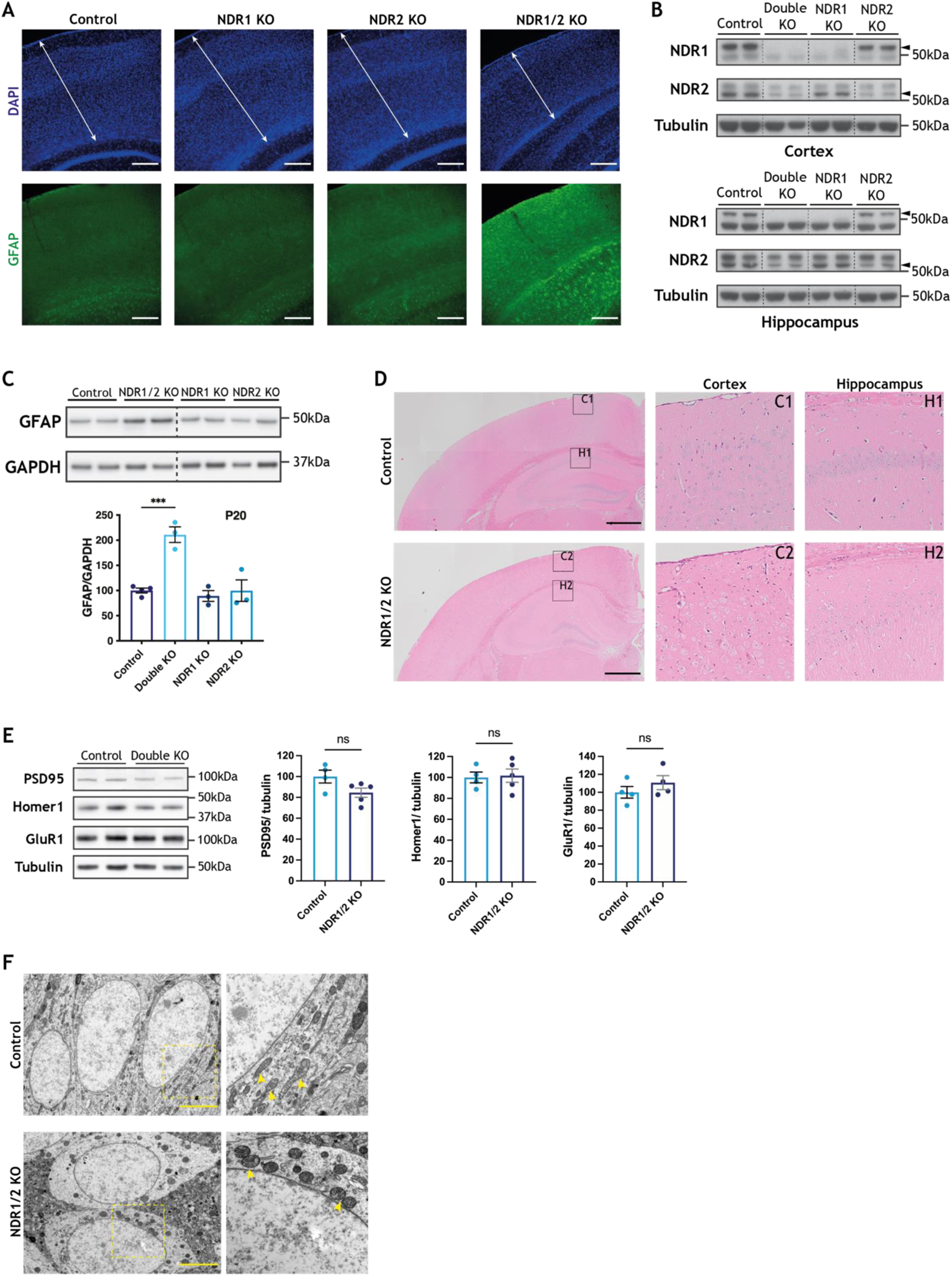
**(A)** Immunofluorescence staining of GFAP in the brains of 20-week-old NDR1 KO, NDR2 KO, NDR1/2 KO and control mice. White arrowed lines indicate the thickness of the cortex. Scale bars 200μm. **(B)** Western blot analyses of NDR1 and NDR2 in brain lysates of 6-week-old NDR1 KO, NDR2 KO, NDR1/2 KO and control mice. Black arrows indicate the correct NDR1 or NDR2 band. **(C)** Western blot analyses of GFAP levels in lysates from the cortex of P20 mice. GAPDH was used as a loading control. The bar graph shows quantifications of the GFAP bands normalised against the GAPDH levels and the data was analysed using an ordinary one-way ANOVA with Tukey’s post hoc test, n = 3-5 mice/ group. **(D)** Haematoxilin and eosin (H&E) stainings of brain slices from 12-week-old NDR1/2 KO and control mice. The black squares indicate the zoomed areas. Scale bars 1000μm. **(E)** Western blot analyses of the synaptic markers PSD95, Homer1 and GluR1 in cortex lysates from 6-week-old mice. The bar graphs show quantifications of PSD95, Homer1 and GluR1 levels normalised against the levels of the loading control tubulin. The data was analysed using unpaired Student’s *t* tests. n=4-5 mice/ group. **(G)** TEM (transmission electron microscope) images from the CA1 area of 12-week-old control and NDR1/2 KO mice. The yellow dotted squares indicate where the higher magnification (9900X) images correspond on the lower magnification (2550X) ones and yellow arrows point out mitochondria. Scale bars 5μm.

**Figure S2.**
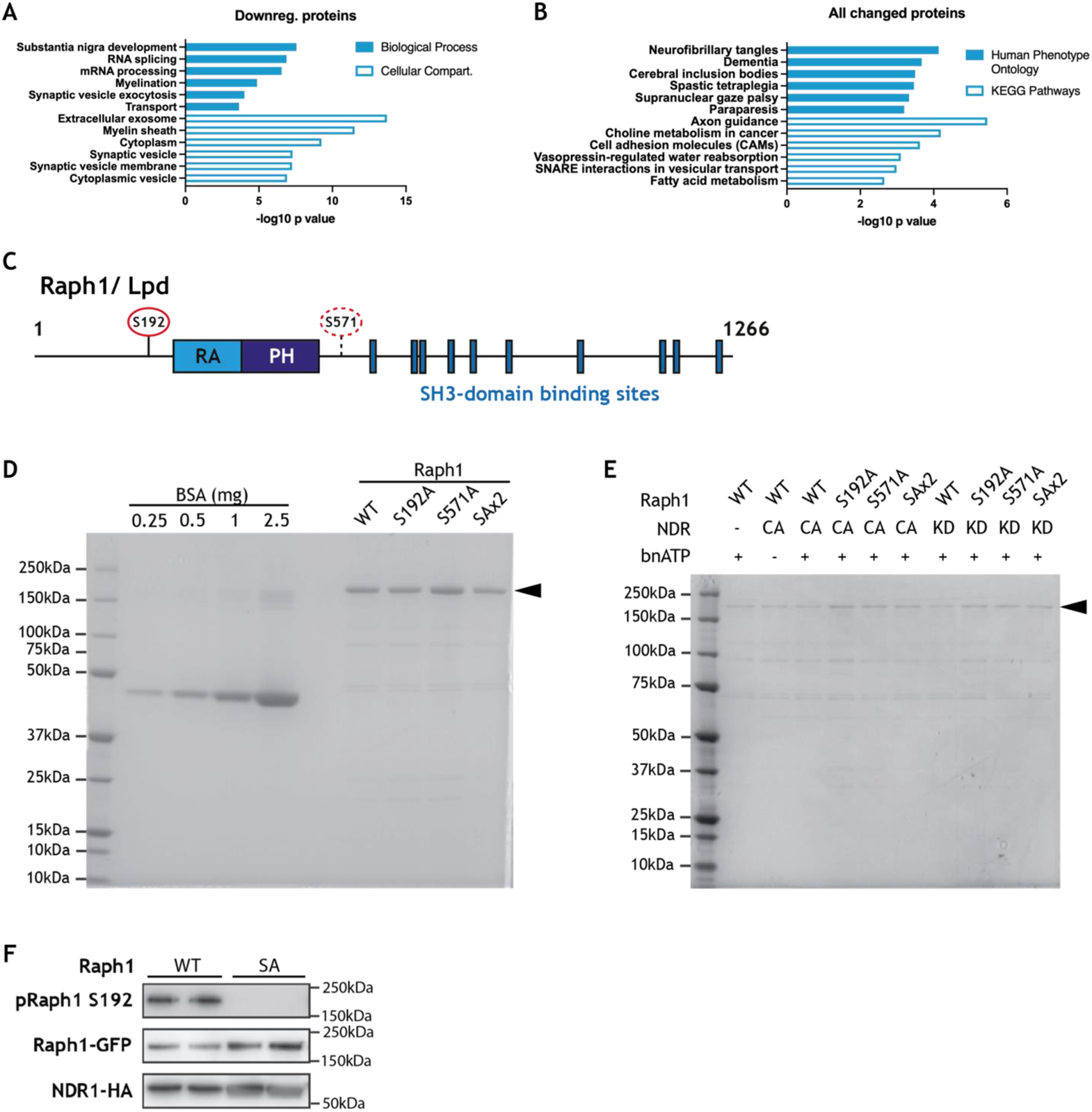
**(A-B)** Results from gene enrichment analyses run using the online tool Database for Annotation, Visualization and Integrated Discovery (DAVID), with the list of genes corresponding to all significantly changed proteins (A) or all significantly downregulated proteins (B) in the proteomics analysis of NDR1/2 KO and control hippocampi. The x axis shows the p-value or EASE score generated by DAVID to show how enriched a term associated with a list of genes is. The top 6 most enriched terms from each category are represented. **(C)** Schematic representation of the structure of mouse Raph1. RA – Ras-associated domain, PH – plekstrin homology domain. S192 is the phosphorylation site targeted by NDR kinases and identified with TMT labelling mass spectrometry. S571 is a putative phosphorylation site which also has the NDR consensus motif. **(D-E)** Coomassie stained gels loaded with samples from purification of Raph1 constructs (D) or from an *in vitro* kinase assay (E). In both images the black arrows show the Raph1 bands. **(F)** Western blot analyses of Raph1 phospho-Ser192, GFP and HA levels in lysates from HEK293T cells transfected with constitutively active NDR1-HA and either wild-type (WT) EGFP-Raph1 or phospho-mutant EGFP-Raph1 S192A (SA).

**Figure S3.**
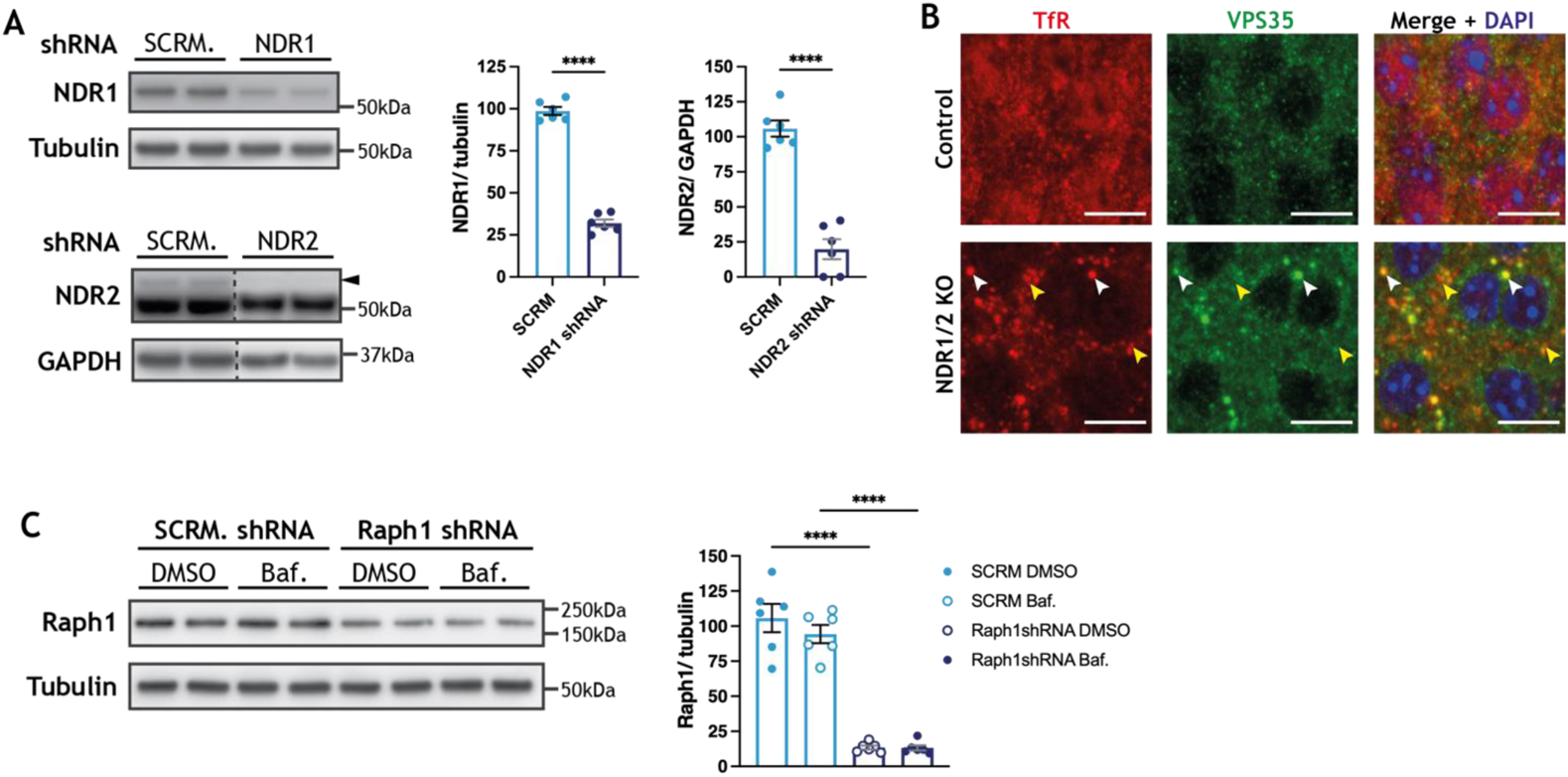
**(A)** Western blot analyses of NDR1 and NDR2 protein levels in lysates from DIV13 rat cortex neurons infected with lentiviral vectors expressing the indicated shRNAs for approximately 5 days. The black arrow shows the correct NDR2 band. The bar graphs show quantifications of the NDR1 and NDR2 bands normalised against the levels of the loading controls. The data was analysed using unpaired Student’s *t* tests. n=6 samples/ group from 3 independent experiments. **(B)** Immunofluorescence stainings of transferrin receptor (TfR) and the retromer component VPS35 in the CA1 area in brain slices from 12-week-old mice. White arrows show co-localisation between TfR and VPS35 and the yellow arrows show TfR puncta that do not co-localise with VPS35. Scale bars 10μm. **(C)** Western blot analyses of Raph1 levels in lysates from DIV13 rat cortex neurons infected with lentiviral vectors expressing scramble shRNA or Raph1 shRNA and treated with either DMSO or 100nM Bafilomycin A1 for 4h. Tubulin was used as a loading control. The bar graph shows quantifications of the Raph1 bands normalised against the tubulin levels and the data was analysed using ordinary one-way ANOVA with Tukey’s post hoc test. n=6 samples/ group from 3 independent experiments.

**Figure S4.**
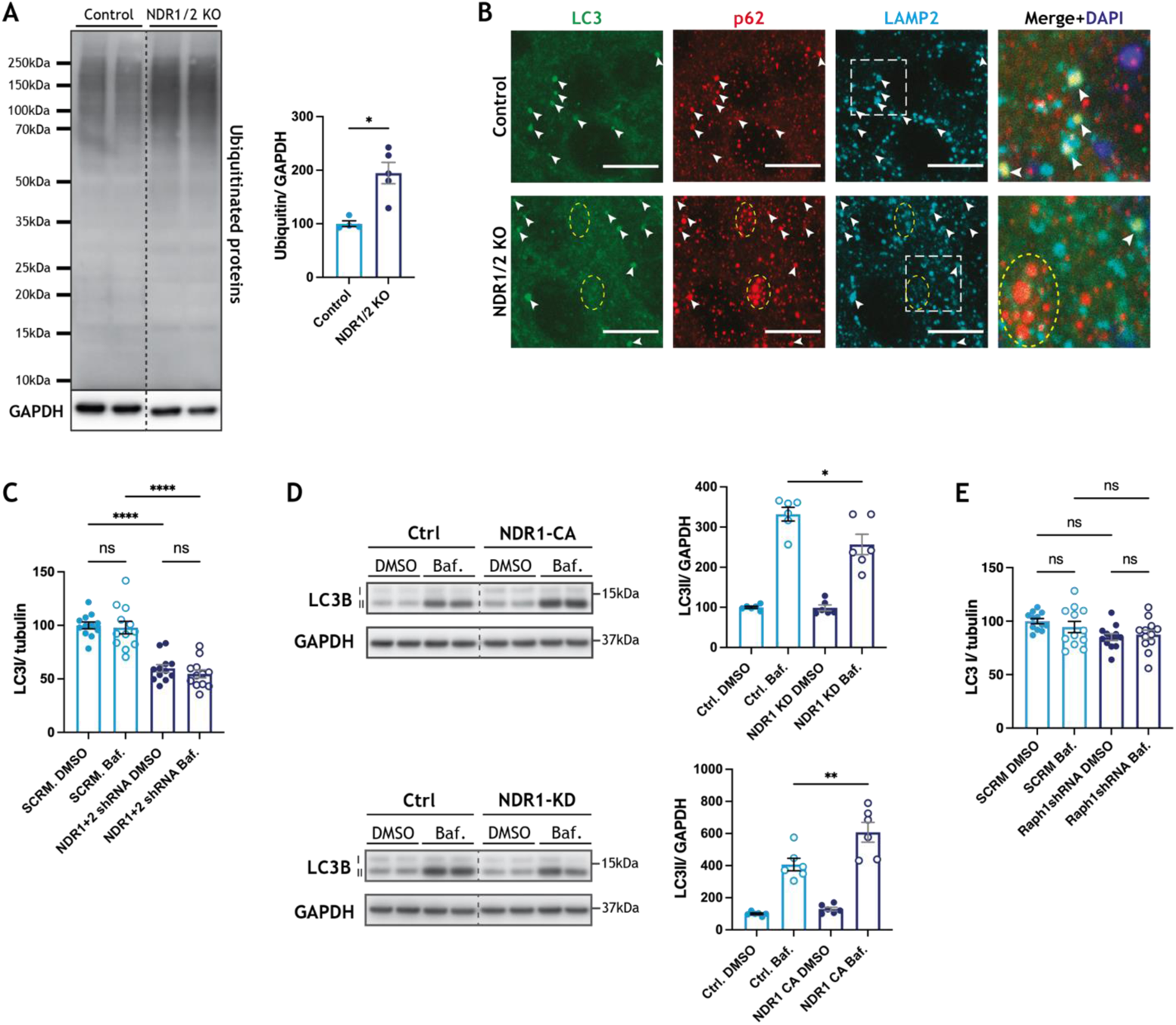
**(A)** Western blot analyses of ubiquitin in lysates from the cortex of 6-week-old NDR1/2 KO and control mice. GAPDH was used as a loading control. The bar graph shows quantifications of the ubiquitin signal within each lane normalised against the GAPDH levels. The data was analysed using an ordinary one-way ANOVA with Tukey’s post hoc test, n = 4-5 mice/ group. **(B)** Western blot analyses of LC3 levels in lysates from HEK293T cells transfected with a constitutively active (CA) NDR1-HA construct, a kinase-dead (KD) NDR1-HA construct or a control HA-expressing plasmid (Ctrl) and treated with either DMSO or 100nM of Bafilomycin A1 (Baf.) for 4 hours prior to lysis. GAPDH was used as a loading control. The graphs show quantifications of the LC3II bands normalised against the tubulin levels. The data was analysed using ordinary one-way ANOVA tests with Tukey’s post hoc test. n = 6 samples/ group from 3 independent experiments. **(C, E)** Quantification of LC3I levels normalised against tubulin levels in lysates from DIV13 rat cortex neurons infected with a scramble shRNA and NDR1 and NDR2 shRNAs (C) or Raph1 shRNA (E). Prior to lysis the neurons were treated with DMSO or with 100nM Bafilomycin A1 (Baf.) for 4h. The data was analysed using ordinary one-way ANOVAs with Tukey’s post hoc test. n=6 samples from 3 independent experiments, 2 technical replicates. **(D)** Immunofluorescence staining of p62, LC3 and the lysosomal marker LAMP2 in the CA1 area of the hippocampus at 12 weeks. White arrows show LC3 puncta, all of which co-localise with both p62 and LAMP2 in control as well as in NDR1/2 KO. The white squares highlight the zoomed areas. Yellow circles show p62 puncta in NDR1/2 KO that do not co-localise with either LC3 or LAMP2. Scale bars 10μm.

**Figure S5.**
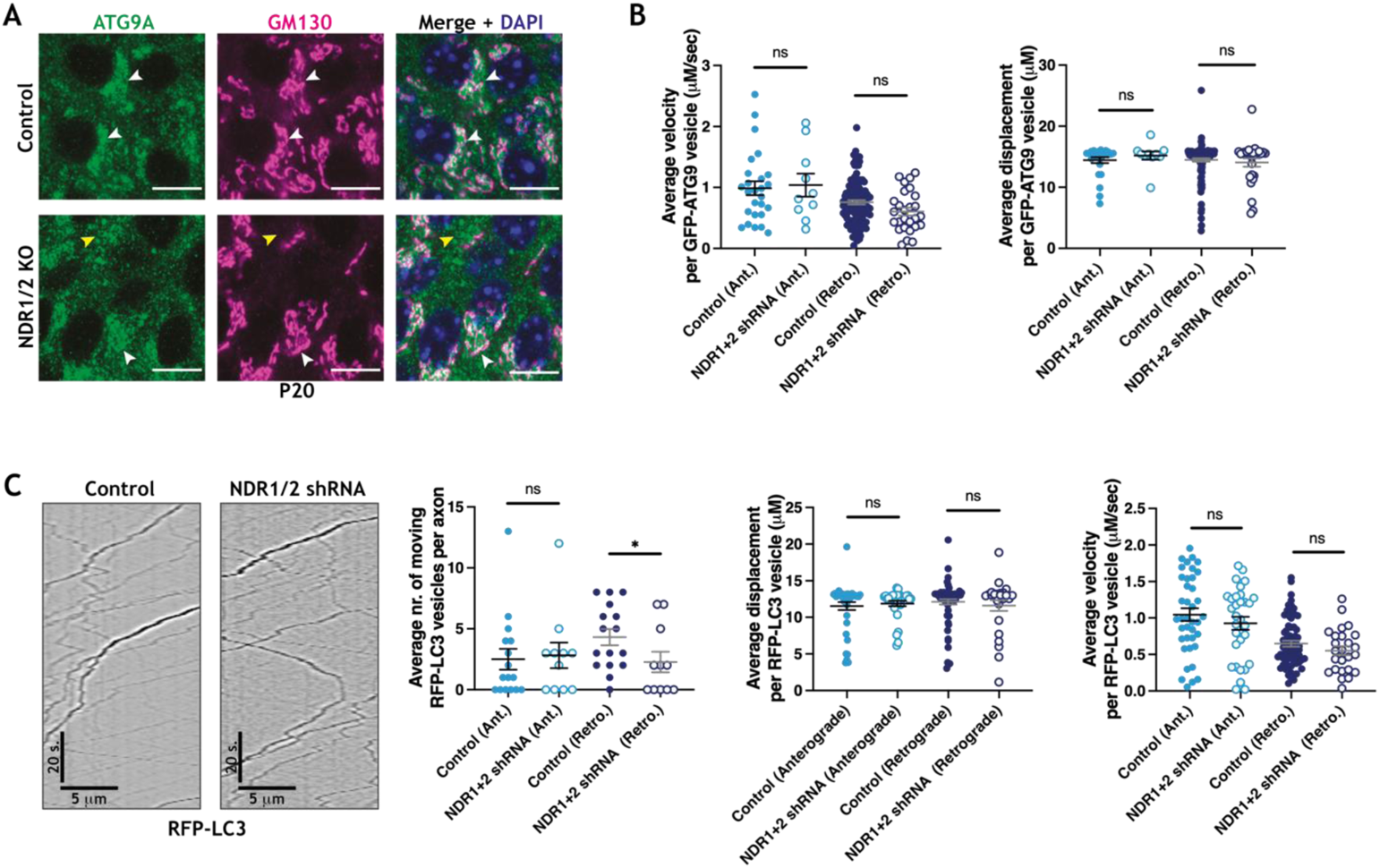
**(A)** Immunofluorescence staining of ATG9A and GM130 in P20 hippocampal brain sections. White arrows mark ATG9A co-localising with GM130 and yellow arrows show ATG9A that does not co-localise with GM130. Scale bars 10μm. **(D)** Analysis of GFP-ATG9A movement in the axons of DIV11 cultured rat hippocampus neurons. n=138 mobile vesicles from 33 cells for scramble shRNA and n=37 mobile vesicles from 24 cells for NDR1/2 shRNA. **(E)** RFP-LC3 live-imaging analysis in axons of DIV11 neurons in culture with representative kymographs (left) and quantifications of RFP-LC3 movement (right - graphs). The cells had been previously infected with a scramble shRNA lentiviral vector as a control or constructs expressing NDR1 and NDR2 shRNAs. Displacement, velocity and total number of moving particles were quantified from kymographs. The data was analysed using Mann-Whitney or unpaired Student’s *t* tests. n=109 mobile particles from 16 cells for scramble shRNA and n=56 mobile particles from 12 cells for NDR1/2 shRNA. The analysed cells were obtained from 3 independent experiments.

## Movie Legends

**Movies 1 and 2**

Confocal image z-stacks of control (Movie 1) and NDR1/2 KO (Movie 2) brain slices stained for ATG9A (green) and p62 (magenta). The stacks are acquired in the hippocampus area, showing the CA1 cell body layer (CA1 c. b.) and stratum oriens (s. o.). The image stack movies start with ATG9A channel, followed by p62 channel and ending with images in which the two channels are merged and DAPI is used to stain nuclei.

**Movies 3 and 4**

Time-lapse imaging of GFP-ATG9A transfected hippocampal primary neurons at DIV11. Neurons were previously infected with lentivirus expressing scrambled shRNA as control (Movie 3) or lentiviruses expressing NDR1 and NDR2 shRNA (Movie 4). The movies are 2 min long and the images are captured with a 500ms interval.

## Supplemental Spreadsheet Legends

**Supplemental Data S1**

Table with all the peptides identified in the TMT labelling mass spectrometry assessment of the total proteome of NDR1/2 KO and control mice and the result of data analysis identifying significantly changed proteins in NDR1/2 KO hippocampi compared to controls.

**Supplemental Data S2**

Tables with the results of gene enrichments analyses conducted with the genes corresponding to upregulated proteins in NDR1/2 KO hippocampi compared to controls.

**Supplemental Data S3**

Tables with the results of gene enrichments analyses conducted with the genes corresponding to downregulated proteins in NDR1/2 KO hippocampi compared to controls.

**Supplemental Data S4**

Tables with the results of gene enrichments analyses conducted with the genes corresponding to all changed proteins in NDR1/2 KO hippocampi compared to controls.

**Supplemental Data S5**

Table with all phosphopeptides identified in the TMT labelling mass spectrometry assessment of the phospho-proteome of NDR1/2 KO and control mice and the result of data analysis identifying significantly changed phosphorylation sites in NDR1/2 KO hippocampi compared to controls.

**Supplemental Data S6**

Tables with the results of gene enrichments analyses conducted with the genes corresponding to all changed phosphorylation sites in NDR1/2 KO hippocampi compared to controls.

